# JUMPlion improves quantitative DIA proteomics through ion-level recovery of missing values

**DOI:** 10.64898/2026.04.28.720212

**Authors:** Yingxue Fu, Zuo-Fei Yuan, Stephanie D. Byrum, Long Wu, Junmin Peng, Xusheng Wang, Anthony A. High

## Abstract

Incomplete quantification remains a persistent challenge in data-independent acquisition (DIA) mass spectrometry (MS), particularly in low-input and single-cell analyses. In identification-driven workflows, missing protein quantities often arise not from true absence of the corresponding peptides, but from failure to retain low-abundance signals from precursor or product ions for quantification. Here we present JUMPlion (local inference of ion-level missingness), a DIA quantification framework that re-examines MS raw files to recover missing values at the ion level before protein quantification. JUMPlion re-extracts precursor- and production signals directly from raw data, infers ion-level measurements within precursor-specific local quantitative neighborhoods, and combines complementary precursor- and production signals into downstream quantification. Using benchmark datasets acquired on multiple DIA platforms, JUMPlion increased protein-level completeness, improved fold-change accuracy, and enhanced detection of differentially abundant proteins while maintaining low differential-abundance false discovery rates. These gains were most evident in low-input and single-cell DIA datasets. Together, these results show that addressing missingness at the ion level before protein-level summarization can improve DIA quantification in diverse acquisition settings.

## Introduction

Data-independent acquisition (DIA) proteomics has emerged as a powerful approach for reproducible proteome-wide quantification in large sample cohorts^1–5^. Relative to data-dependent acquisition (DDA), DIA offers improved sampling consistency and greater robustness for comparative proteomics, making it particularly well suited for studies requiring broad proteome coverage in many samples^6–8^. Despite these advantages, incomplete quantitative measurements remain prevalent in DIA datasets and can bias fold-change estimation, reduce statistical power, and complicate downstream analyses^9–12^. This challenge is especially pronounced in low-input and single-cell proteomics experiments, where reduced signal intensity increases variability in the detection of precursor- and production signals across runs^13–15^.

In current DIA workflows, precursor- and protein-level quantities are typically derived from production intensities through precursor-level or direct protein-level summarization^16–19^. Aggregation of production measurements can reduce apparent missingness at the precursor and protein levels^20, 21^. Accordingly, missing values in DIA analyses are commonly addressed using post hoc imputation applied to peptide- or protein-level matrices after summarization^22–24^. However, missingness at these higher levels often originates earlier in the workflow, when low-abundance ion signals are not retained for downstream quantification during identification-driven processing. Because identification-driven DIA algorithms generally require precursor-level evidence to exceed confidence thresholds for false discovery rate (FDR) control, measurable but low-intensity ion signals may yield sub-threshold identification scores and thus be excluded from quantification, even when quantitative evidence is present in the raw data and is technically extractable^25^. Consequently, missing quantitative values often reflect incomplete signal observability rather than the true biological absence of the precursor. These considerations suggest that missingness in DIA should not be viewed solely as a matrix-level imputation problem after summarization, but also as an ion-level signal observability problem that may be addressed directly in the raw data. In addition, precursor signals from MS1 spectra are often not systematically incorporated into final quantitative estimates, despite their potential to provide complementary evidence for precursor abundance^26, 27^. As a result, informative signal present in the raw data may remain underutilized.

Here, we present JUMPlion, a DIA quantification framework designed to recover precursor-associated quantitative evidence prior to precursor- and protein-level summarization (**Fig. 1**). JUMPlion re-extracts precursor and product ion signals directly from raw data, infers missing ion-level measurements within precursor-specific local quantitative neighborhoods, and supports the integration of complementary MS1- and MS2-derived evidence into downstream quantification. By operating at the ion level rather than only after higher-level aggregation, JUMPlion aims to recover measurable low-abundance signals that may have been excluded during identification-driven processing. We evaluated the performance of JUMPlion using benchmark datasets acquired on multiple DIA platforms from different vendors, including low-input and single-cell DIA datasets.

**Fig. 1.**
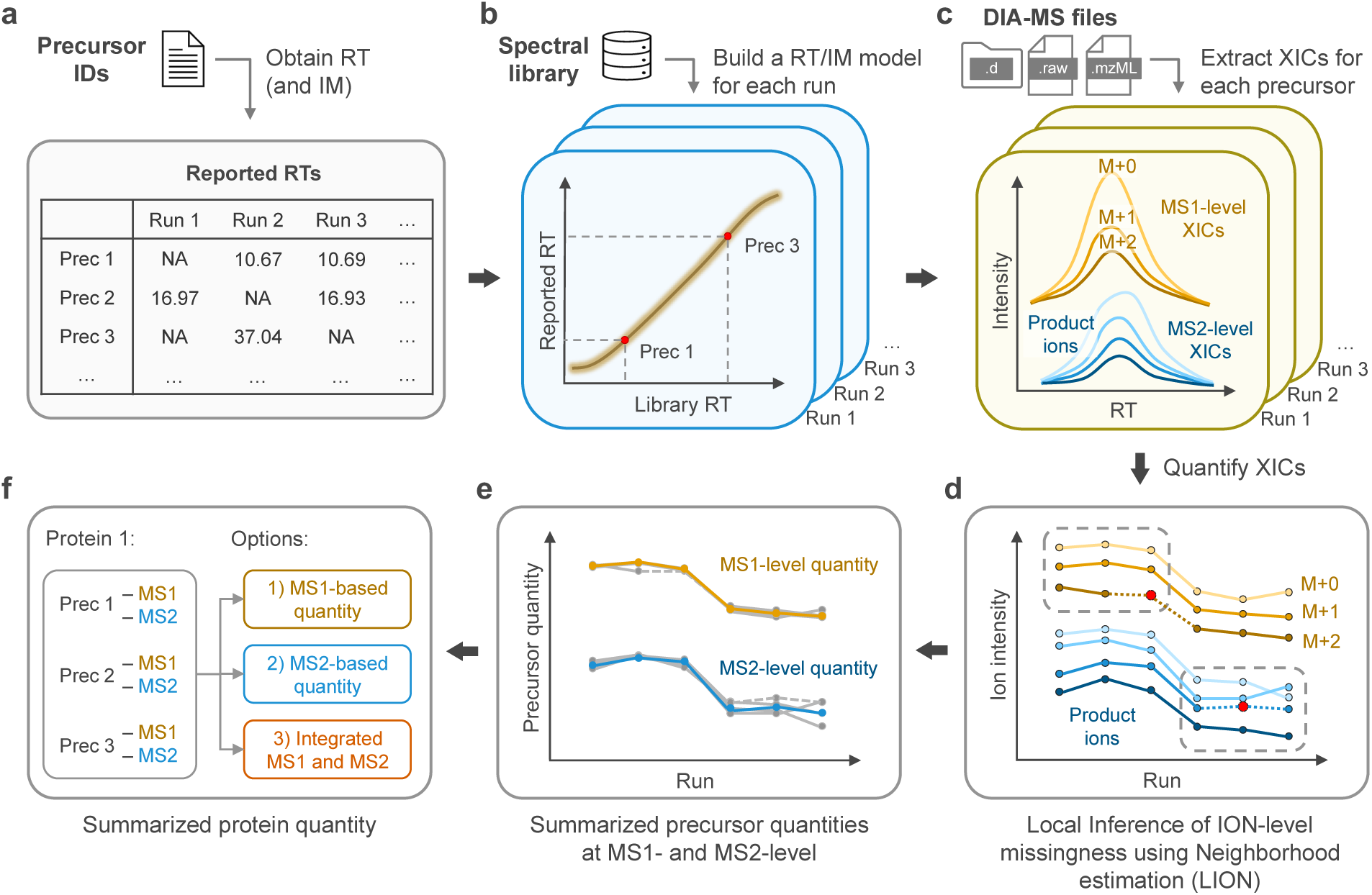
Schematic overview of the JUMPlion workflow. **a,b**, JUMPlion extracts reported retention times (RTs) from precursor-level identification output and builds run-specific RT prediction models by integrating these values with indexed retention times (iRTs) from the spectral library. For timsTOF datasets, ion mobility (IM) prediction models are constructed in parallel. Precursors lacking reported RT or IM values are assigned predicted values from these models. **c**, JUMPlion retrieves extracted ion chromatograms (XICs) directly from raw data for all identified precursors at both MS1 and MS2 levels. MS1-level XICs are extracted for the monoisotopic peak and predicted observable isotopic peaks, whereas MS2-level XICs are extracted for the top six product ions defined in the spectral library. **d**, XICs are quantified in each run, and ion-level intensities are organized into precursor-specific cross-run ion intensity vectors. Missing values within MS1- and MS2-derived vectors are inferred independently using the Local Inference of ION-level missingness using Neighborhood estimation (LION) strategy within condition-specific neighborhoods. **e**, MS1- and MS2-derived precursor quantities are summarized from cross-run ion intensity vectors after LION-based inference. **f**, Protein-level quantities are summarized using MS1-only, MS2-only, or integrated MS1/MS2 strategies to generate a complete quantification matrix without missing values.

## Results

### Ion-level missingness is extensive in DIA and can be recovered to generate more complete protein-level quantification

To evaluate the extent of missingness at different levels of quantification, we analyzed three benchmark datasets acquired on timsTOF Pro 2, Exploris 480, and Orbitrap Astral instruments, each comprising defined mixtures of human, yeast, and *E. coli* peptides at known proportions^28,29^ (**Supplementary Table 1**). In all three datasets, missingness was greatest at the production level and progressively reduced after summarization to precursors and proteins (**Fig. 2a** and **Supplementary Fig. 1a,e**). For example, in the Orbitrap Astral dataset, only 29% of product ions were quantified in every run, compared with 53% of precursors and 83% of protein groups (**Supplementary Fig. 1e**). Production missingness was strongly associated with lower signal intensity, whereas this relationship was attenuated following summarization to precursor- and protein-level quantities (**Fig. 2b–d** and **Supplementary Fig. 1**). These observations indicate that higher-level summarization reduces the apparent extent of missingness while obscuring its relationship to the underlying ion-level signal intensity.

**Fig. 2.**
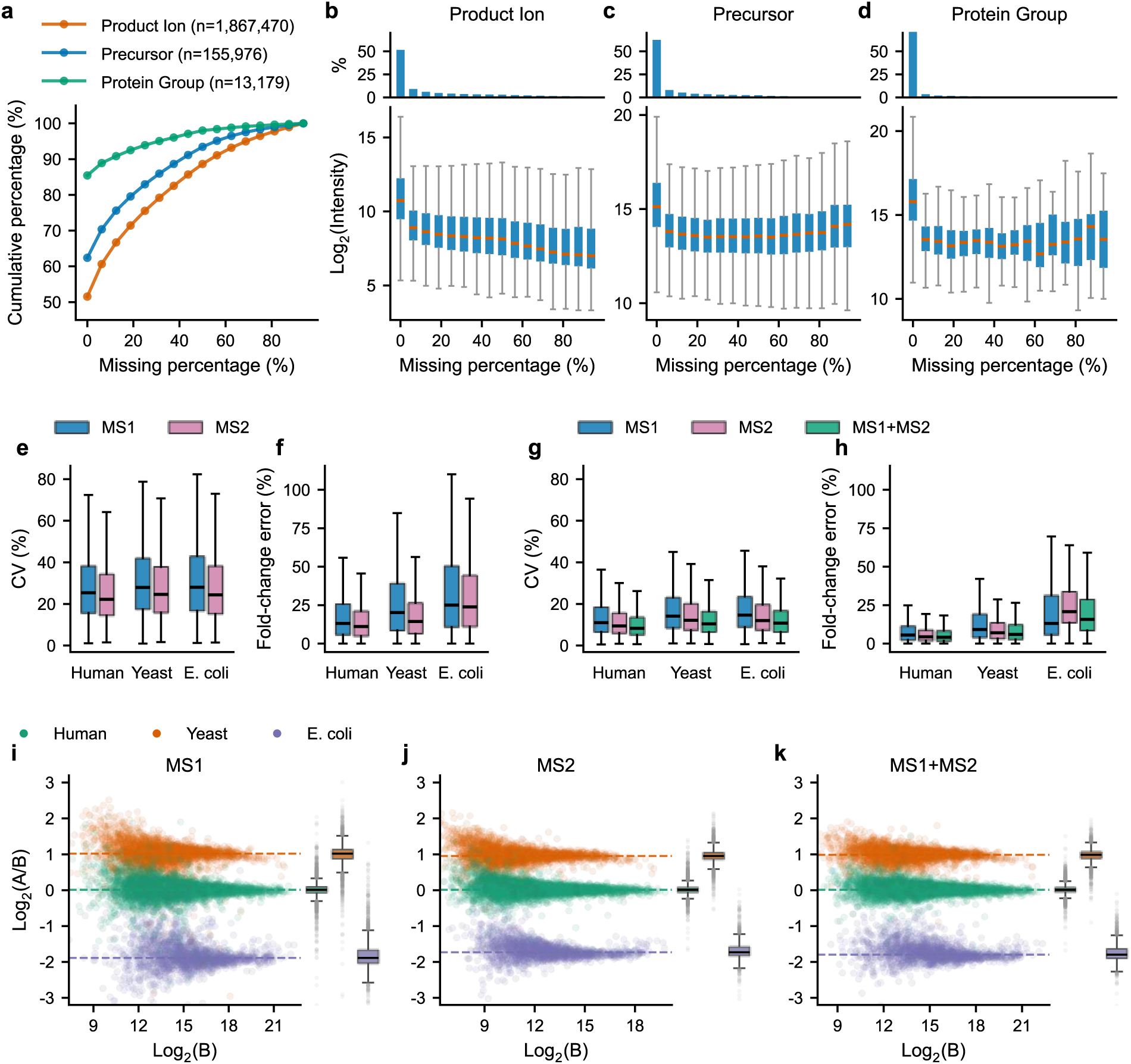
JUMPlion generates complete protein-level quantification profiles for DIA proteomics. **a**, Cumulative percentages of product ions, precursors, and protein groups on different levels of missingness. **b–d**, Relationship between missing-value percentage and log_2_ intensity at the production (**b**), precursor (**c**), and protein-group (**d**) levels. **e,f**, Coefficient of variation (CV; **e**) and fold-change error (**f**) of precursor quantification summarized from MS1- and MS2-level ion intensities. **g,h**, CV (**g**) and fold-change error (**h**) of protein quantities summarized using MS1-only, MS2-only, and integrated MS1/MS2 strategies. **i–k**, Scatter plots of log_2_-transformed protein ratios versus log_2_-transformed protein intensities and corresponding box plots of log_2_-transformed ratios. Colored dashed lines indicate median log_2_(A/B) values for human (green), yeast (orange), and *E. coli* (purple) proteins. Panels **i**, **j**, and **k** show protein quantities summarized using MS1-only, MS2-only, and integrated MS1/MS2 strategies, respectively. Results for the timsTOF Pro 2 (400 ng) dataset are shown.

To recover quantitative signals that may be excluded during identification-driven processing, JUMPlion re-extracts precursor- and production signals directly from raw data. Signal extraction was performed for precursors passing run-specific identification FDR (default: < 0.01) and identified in multiple runs (default: ≥ 3). For precursors lacking reported chromatographic coordinates, JUMPlion inferred run-specific retention time (RT) values using models trained from observed identifications and spectral-library indexed retention time (iRT) information (**Fig. 1a,b**). For timsTOF datasets, ion mobility (IM) values were inferred in parallel. In the three benchmark datasets, inferred RT/IM values were required for approximately 10–17% of precursor coordinates (timsTOF Pro 2, 10.44%; Exploris 480, 16.87%; Orbitrap Astral, 13.91%) (**Supplementary Fig. 2**). Using observed or inferred RT/IM coordinates, JUMPlion retrieved extracted ion chromatograms (XICs) for precursor ions from MS1 spectra and for product ions from MS2 spectra (**Fig. 1c**). In contrast to conventional DIA quantification workflows, which rely predominantly on production measurements for downstream summarization^16–18^, JUMPlion additionally captures MS1-level XICs corresponding to monoisotopic precursor ions and predicted isotopic peaks. The number of MS1-level cross-run ion intensity vectors increased with precursor peptide mass. In the three datasets, 72–98% of precursors were represented by at least three such vectors (timsTOF Pro 2, 98%; Exploris 480, 78%; Orbitrap Astral, 72%) (**Fig. 1d** and **Supplementary Fig. 3)**.

Missing ion intensities were then inferred separately at the MS1 and MS2 levels using a procedure termed Local Inference of ION-level missingness using Neighborhood estimation (LION) (**Fig. 1d**). For each precursor, missing ion values were estimated from neighborhood-specific minimum intensities within samples sharing the same biological condition. Because ion-level missingness was associated with reduced signal intensity (**Fig. 2b** and **Supplementary Fig. 1b,f**), the local minimum XIC intensity provided a natural estimate for incompletely observed signals. Evaluation of scaling factors applied to this local minimum showed that coefficients of variation (CVs) decreased with increasing scale before reaching a plateau, whereas deviation from expected fold change increased monotonically (**Supplementary Fig. 4 and 5**). A scaling factor of 1.0 was therefore selected as a balance between quantitative variability and fold-change accuracy for all three acquisition platforms.

We next evaluated the quantitative performance of precursor-level measurements summarized from LION-inferred MS1- and MS2-derived ion intensities (**Fig. 1e**). In all three benchmark datasets, precursor quantities summarized from MS2 ions showed lower CVs and smaller fold-change errors than those summarized from MS1 ions, with the differences most pronounced in the Exploris 480 and Orbitrap Astral datasets (**Fig. 2e,f** and **Supplementary Fig. 6**). These precursor quantities were then summarized to the protein level using three strategies: MS1-only, MS2-only, or integrated MS1/MS2 precursor quantities (**Fig. 1f**). All three strategies produced complete protein-level quantification matrices with no missing values. In the timsTOF Pro 2 dataset, integrating MS1- and MS2-derived precursor quantities reduced both CV and fold-change error relative to either MS1-only or MS2-only summarization (**Fig. 2g–k**). In contrast, in the Exploris 480 and Orbitrap Astral datasets, integrated MS1/MS2 summarization performed similarly to, or slightly worse than, MS2-only quantification (**Supplementary Fig. 6**). Together, these results indicate that ion-level signal recovery can generate complete protein-level quantification profiles on different DIA platforms, whereas the benefit of incorporating MS1 evidence depends on acquisition-specific sampling characteristics.

### JUMPlion increases detection of differentially abundant proteins while maintaining false discovery rate control

We next asked whether recovering ion-level signals before summarization improves downstream differential abundance analysis. To this end, we used the benchmark mixtures to define true positive (TP) and false positive (FP) differentially abundant proteins (DAPs) based on known species-specific abundance differences, and calculated differential abundance FDR (daFDR) as FP/(FP + TP)^30^.

Before benchmarking JUMPlion against existing DIA quantification workflows, we first determined which protein-level summarization strategy performed best on each platform. In the timsTOF Pro 2 dataset, integration of MS1- and MS2-derived precursor quantities yielded the highest number of TP DAPs (5,058), compared with MS1-only (4,811) and MS2-only (4,995) quantification, and also achieved the lowest daFDR (1.02% versus 3.66% for MS1-only and 1.44% for MS2-only) (**Supplementary Fig. 7a–c**). In the Exploris 480 dataset, MS2-only and integrated MS1/MS2 quantification performed similarly, with 3,786 and 3,800 TP DAPs and daFDR values of 1.66% and 1.76%, respectively (**Supplementary Fig. 7d–f**). In the Orbitrap Astral dataset, MS1-only quantification performed substantially worse (3,090 TP DAPs; daFDR 24.95%) than MS2-only quantification (5,221 TP DAPs; daFDR 5.26%) (**Supplementary Fig. 7g–i**). Based on these results, integrated MS1/MS2 quantification was used for timsTOF Pro 2, whereas MS2-based quantification was used for Exploris 480 and Orbitrap Astral in subsequent benchmarking.

We then compared JUMPlion (v0.8.6) with DIA-NN^17^ (v2.2) and Spectronaut^2^ (v20.3). In all three benchmark datasets, JUMPlion yielded a larger number of quantifiable proteins than either DIA-NN or Spectronaut (**Supplementary Fig. 8**). In the Orbitrap Astral dataset, for example, JUMPlion quantified 14,450 proteins, compared with 13,430 and 13,281 proteins using DIA-NN and Spectronaut, respectively (**Supplementary Fig. 8g–i**). Relative to DIA-NN- and Spectronaut-derived protein quantities, JUMPlion-derived measurements showed modestly increased CVs but lower fold-change error (**Supplementary Fig. 8**), indicating that the additional quantitative signal recovered by JUMPlion preserved expected abundance differences more accurately despite somewhat greater dispersion.

This improvement translated into increased sensitivity for differential abundance detection (**Fig. 3** and **Supplementary Fig. 9 and 10**). In the timsTOF Pro 2 dataset, for example, JUMPlion identified 5,058 TP DAPs at a daFDR of 1.02%, compared with 4,715 TPs at 0.46% daFDR for DIA-NN and 4,556 TPs at 0.63% daFDR for Spectronaut (**Fig. 3a–c**). Most DAPs detected using JUMPlion overlapped with those identified by DIA-NN and/or Spectronaut (e.g., 96.3% of yeast proteins and 92.2% of *E. coli* proteins for the timsTOF Pro 2 dataset) (**Fig. 3d,e** and **Supplementary Fig. 9d,e and 10d,e**). Notably, proteins identified as differentially abundant only by JUMPlion were biased toward lower abundance relative to all JUMPlion-quantified proteins from the same species (**Fig. 3f,g** and **Supplementary Fig. 9f,g and 10f,g**), consistent with preferential rescue of low-abundance signals that are more likely to be lost in identification-driven workflows. Receiver operating characteristic (ROC) analysis further showed that, in the timsTOF Pro 2 dataset, JUMPlion achieved a partial area under the curve (pAUC) comparable to DIA-NN (0.0988 versus 0.0991), whereas in the Exploris 480 and Orbitrap Astral datasets DIA-NN showed the highest pAUC, followed by JUMPlion and Spectronaut (**Supplementary Fig. 11**). Together, these results show that JUMPlion improves recovery of true differential protein abundance signals while maintaining empirical false discovery rate control.

**Fig. 3.**
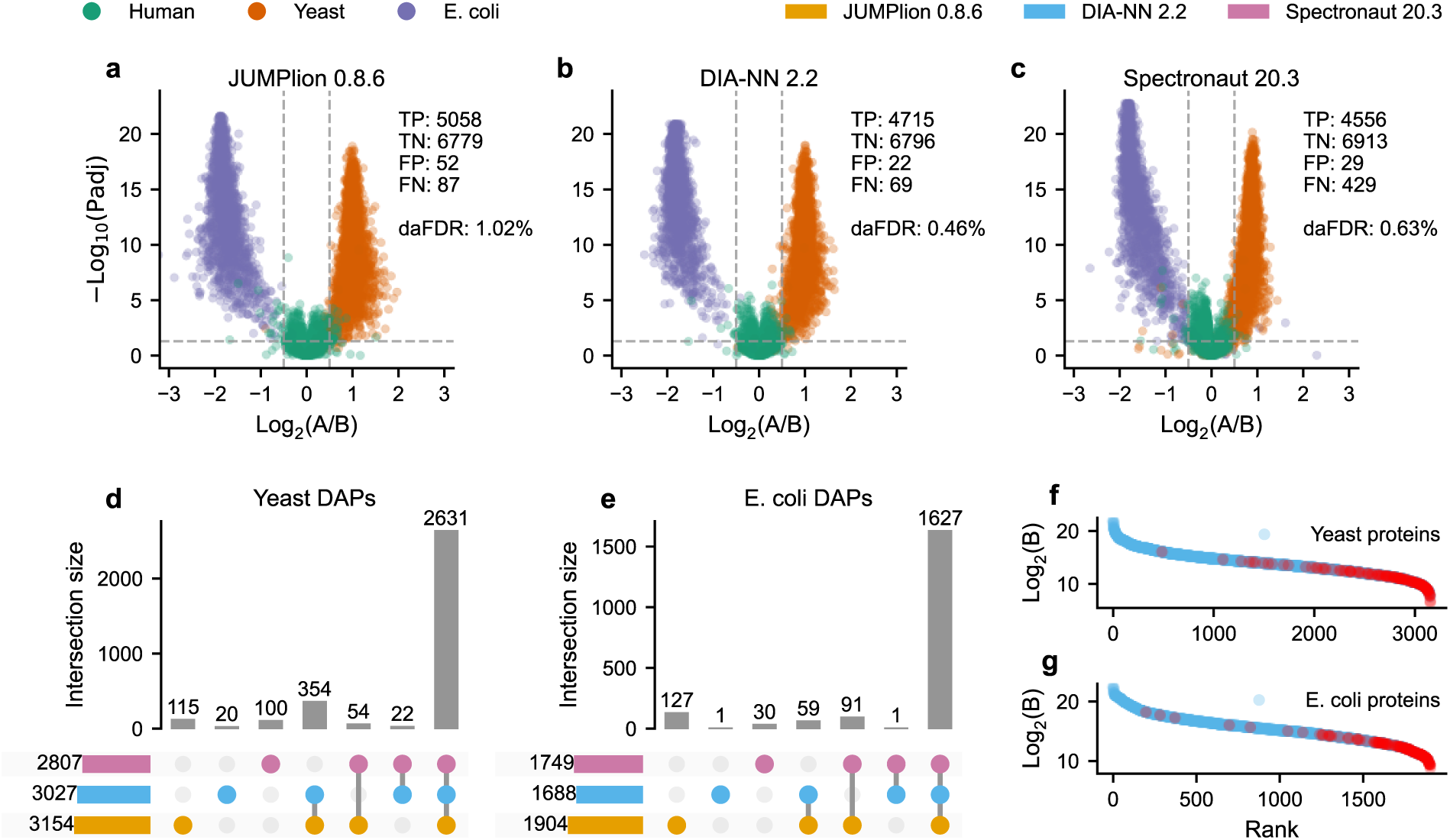
JUMPlion increases detection of differentially abundant proteins while maintaining false discovery rate control. **a–c**, Volcano plots of differential protein abundance analysis using protein quantities derived from JUMPlion v0.8.6 (**a**), DIA-NN v2.2 (**b**), and Spectronaut v20.3 (**c**). Points are colored by species of origin. The numbers of true positives (TP), true negatives (TN), false positives (FP), and false negatives (FN), together with the differential abundance false discovery rate (daFDR; FP/(TP+FP)), are indicated. **d,e**, UpSet plots showing unique and shared differentially abundant proteins (DAPs) from yeast (**d**) and *E. coli* (**e**) identified by JUMPlion, DIA-NN, and Spectronaut. **f,g**, Ranked log_2_ intensities of all yeast (**f**) and *E. coli* (**g**) proteins, with DAPs uniquely identified using JUMPlion-derived quantities highlighted in red. Results for the timsTOF Pro 2 (400 ng) dataset are shown.

### JUMPlion improves differential abundance analysis in low-input DIA datasets

We next assessed whether the advantages of ion-level signal recovery extend to low-input DIA, where reduced signal intensity is expected to increase measurement sparsity. To test this, we analyzed a timsTOF Pro 2 benchmark dataset acquired with a peptide loading amount of 200 pg^31^ and compared it with a standard-loading dataset acquired at 200 ng^28^, representing an approximately three-order-of-magnitude difference in input (**Supplementary Fig. 12a**). As expected, DIA-NN-derived quantification of the 200 pg dataset yielded far fewer quantified precursors (15,836 versus 129,945) and protein groups (3,372 versus 11,661) than the 200 ng dataset. Data completeness was also markedly reduced at low input: only 30.2% of identified precursors and 53.0% of protein groups were quantified in all runs in the 200 pg dataset, compared with 57.5% and 81.8%, respectively, in the 200 ng dataset (**Supplementary Fig. 12b,c**).

We then evaluated JUMPlion-derived protein quantities in the 200 pg dataset using MS1-only, MS2-only, and integrated MS1/MS2 summarization. Integration of MS1- and MS2-derived precursor quantities yielded the best overall performance, increasing the number of TP DAPs while reducing the daFDR and improving pAUC relative to either MS1-only or MS2-only summarization (**Supplementary Fig. 12d–i**). Using this integrated strategy, JUMPlion produced 2,400 quantifiable proteins, compared with 2,043 and 2,601 quantifiable proteins by DIA-NN and Spectronaut, respectively (**Fig. 4a**). Although Spectronaut quantified a larger number of proteins overall, JUMPlion provided lower fold-change error than both DIA-NN and Spectronaut, while CVs were intermediate between the two methods (**Fig. 4b,c**), indicating a favorable balance between quantitative stability and accuracy in this signal-limited setting.

**Fig. 4.**
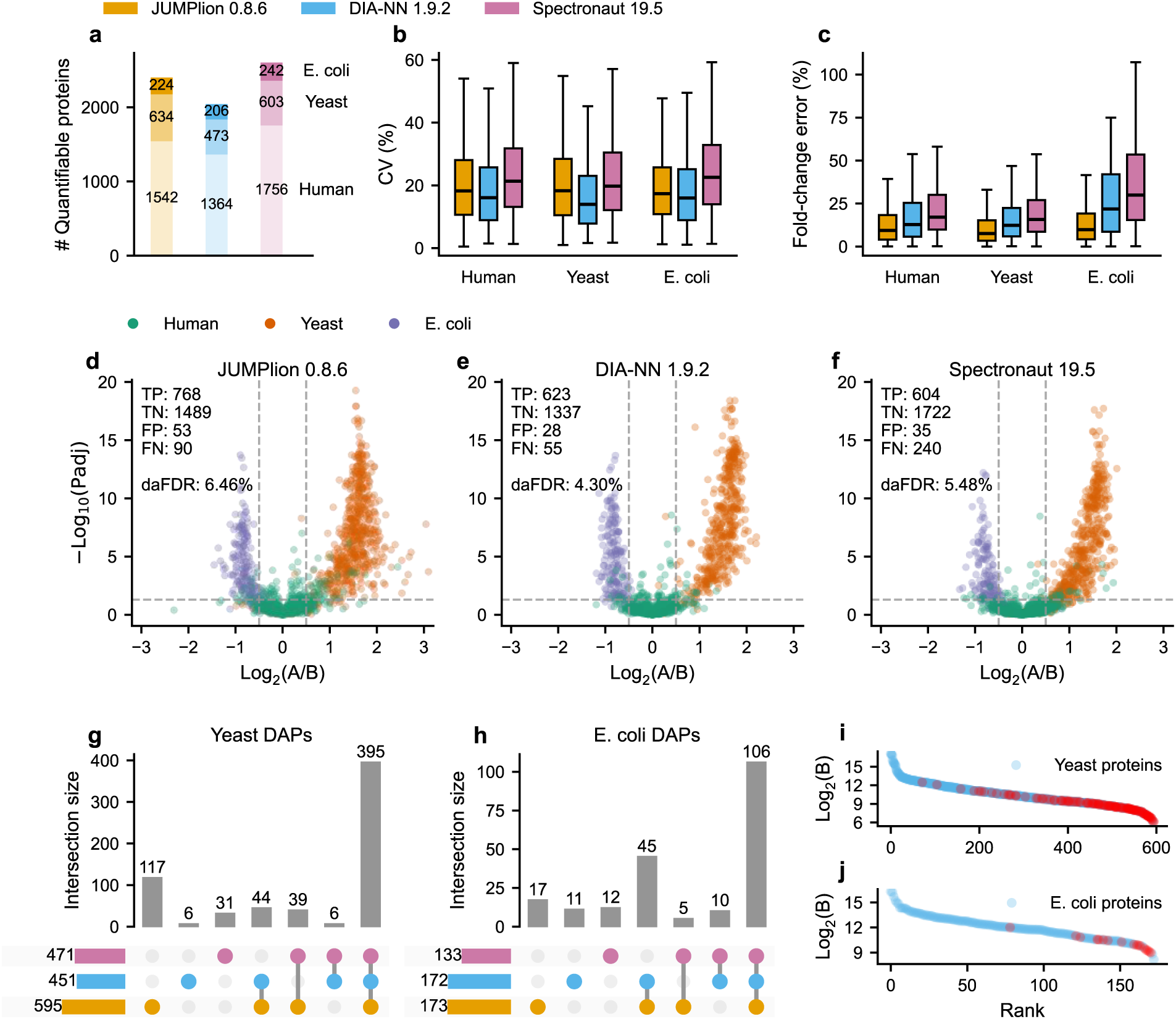
JUMPlion improves differential abundance analysis in low-input DIA datasets. **a–c**, Number of quantifiable proteins (**a**), coefficient of variation (CV; **b**), and fold-change error (**c**) based on protein quantities derived from JUMPlion v0.8.6, DIA-NN v1.9.2, and Spectronaut v19.5. **d–f**, Volcano plots of differential protein abundance analysis using protein quantities derived from JUMPlion v0.8.6 (**d**), DIA-NN v1.9.2 (**e**), and Spectronaut v19.5 (**f**). Points are colored by species of origin. The numbers of TP, TN, FP, and FN, together with dapFDR (FP/(TP+FP)), are indicated. **g,h**, UpSet plots showing unique and shared DAPs from yeast (**g**) and *E. coli* (**h**) identified by JUMPlion, DIA-NN, and Spectronaut. **i,j**, Ranked log_2_ intensities of all yeast (**i**) and *E. coli* (**j**) proteins, with DAPs uniquely identified using JUMPlion-derived quantities highlighted in red.

These gains improved downstream differential abundance analysis. JUMPlion identified 768 TP proteins, compared with 623 and 604 using DIA-NN and Spectronaut, respectively, while maintaining daFDR within a comparable range (**Fig. 4d–f**). Among DAPs detected using JUMPlion, 19.7% of yeast proteins (117 of 595) and 9.8% of *E. coli* proteins (17 of 173) were uniquely identified by JUMPlion (**Fig. 4g,h**). As in the cross-platform benchmarks, proteins uniquely detected as differentially abundant by JUMPlion were shifted toward lower abundance relative to all JUMPlion-quantified proteins within the same species (**Fig. 4i,j**). These results indicate that ion-level recovery is particularly beneficial in low-input DIA, where conventional identification-driven workflows are more likely to lose low-abundance quantitative evidence.

### JUMPlion resolves treatment-associated proteome changes in single-cell DIA data

To determine whether JUMPlion can support downstream biological analysis in highly sparse single-cell DIA data, we applied it to a timsTOF Pro single-cell proteomics dataset in which individual MCF-7 breast cancer cells were treated with doxorubicin or vehicle control (DMSO)^31^. Downstream analysis was restricted to samples containing matched amounts of yeast and *E. coli* spike-in peptides^31^. Under these conditions, JUMPlion generated complete protein-level quantification profiles for 2,937 proteins, whereas DIA-NN- and Spectronaut-derived outputs yielded 1,065 and 1,303 proteins with no missing values, respectively (**Fig. 5a**). When proteins were considered quantifiable if at least five non-missing values were present among the 15 samples in each treatment group, 2,937, 2,678, and 3,197 proteins were quantified using JUMPlion, DIA-NN, and Spectronaut, respectively, of which 2,519 were shared among all three methods (**Fig. 5b**).

**Fig. 5.**
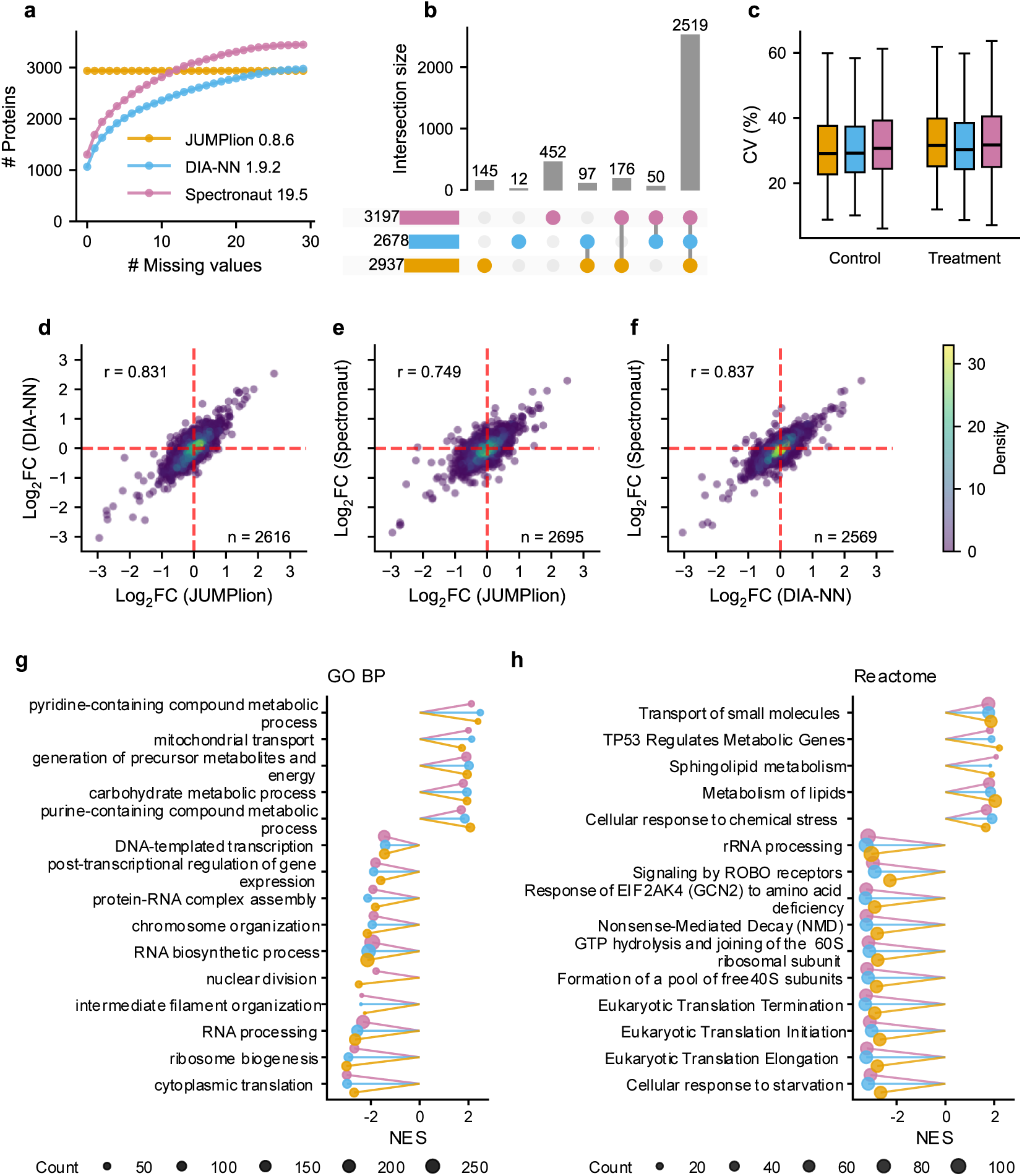
JUMPlion resolves treatment-associated proteome changes in single-cell DIA data. **a**, Cumulative numbers of proteins on different levels of missingness based on protein quantities derived from JUMPlion v0.8.6, DIA-NN v1.9.2, and Spectronaut v19.5. JUMPlion generated a complete protein-level matrix without missing values. **b**, UpSet plot showing unique and shared quantifiable proteins identified by JUMPlion, DIA-NN, and Spectronaut. **c**, Distributions of within-group CVs for control and doxorubicin-treated cells based on protein quantities derived from JUMPlion v0.8.6, DIA-NN v1.9.2, and Spectronaut v19.5. **d–f**, Pairwise correlations of protein-level log_2_FCs from differential abundance analysis: JUMPlion versus DIA-NN (**d**), JUMPlion versus Spectronaut (**e**), and DIA-NN versus Spectronaut (**f**). **g,h**, Normalized enrichment scores (NES) of significantly enriched Gene Ontology Biological Process gene sets (**g**) and Reactome pathways (**h**) identified by gene set enrichment analysis (GSEA) using protein-level log_2_FCs. All displayed pathways were significant at adjusted *P* < 0.05.

To assess whether increased completeness altered global data structure, protein-level quantities generated by the three software tools were normalized using SCnorm^32^ and then batch-corrected using limma^33^ (Methods; **Supplementary Fig. 13a–c**). Principal component analysis of normalized and batch-corrected data showed clear separation between doxorubicin-treated and control cells without batch-driven clustering for any of the three quantification approaches (**Supplementary Fig. 13d–f**). The proportion of variance explained by the first two principal components was similar among methods (36.64% for JUMPlion, 36.53% for DIA-NN, and 39.17% for Spectronaut), and within-group CV distributions were also comparable after normalization and batch correction (**Fig. 5c**). Thus, the increased completeness of JUMPlion-derived profiles did not distort the major biological or technical structure of the data.

We next tested whether this improved quantitative basis enhanced detection of treatment-associated proteome changes. Using thresholds of |log_2_FC| > log_2_1.5 and adjusted *P* < 0.05, differential abundance analysis identified 240 proteins with significant doxorubicin-associated abundance changes using JUMPlion-derived quantities, compared with 195 and 160 proteins using DIA-NN and Spectronaut, respectively (**Supplementary Fig. 13g–i**). Protein-level log_2_FC values were highly concordant between JUMPlion and DIA-NN (*r* = 0.831), similar to the agreement observed between DIA-NN and Spectronaut (**Fig. 5d–f**). Gene set enrichment analysis^34^ using Gene Ontology Biological Process^35^ and Reactome^36^ gene sets recovered similar top positively and negatively enriched pathways among all three quantification approaches (**Fig. 5g,h**), including TP53 Regulates Metabolic Genes, cellular response to chemical stress, and mitochondrial transport among positively enriched pathways, and DNA-templated transcription, RNA processing, and eukaryotic translation initiation among negatively enriched pathways. Together, these data show that JUMPlion improves quantitative completeness in single-cell DIA while preserving biologically coherent treatment-associated signals and enabling sensitive downstream analysis.

## Discussion

Incomplete quantitative measurements remain a persistent challenge in DIA proteomics, particularly in low-input and single-cell applications, where weak signal and variable ion observability contribute to missingness across runs^13^. Here, we show that a substantial fraction of this missingness can be addressed before precursor- and protein-level summarization by operating directly at the ion level. Rather than viewing missing values primarily as a downstream matrix-level problem, JUMPlion treats them, in many cases, as a consequence of incomplete retention or observability of precursor-associated ion signals during identification-driven processing. By re-extracting precursor- and production evidence from raw data and applying local ion-level inference before summarization, JUMPlion improves quantitative completeness and enhances sensitivity for downstream differential abundance analysis in multiple DIA settings.

This strategy differs from conventional post hoc matrix-level imputation, which is typically applied after peptide- or protein-level summarization^24^. By that stage, precursor-specific ion relationships have already been compressed into aggregate estimates^37^. JUMPlion instead intervenes earlier by directly recovering measurable MS1- and MS2-derived ion signals, and then inferring values that remain missing in a local, precursor-specific manner. Because ion detectability is shaped by multiple precursor-specific factors, including abundance, chromatographic behavior, co-elution, and acquisition context, local inference helps preserve precursor-specific signal structure while limiting inappropriate borrowing from unrelated ions. In addition, inference is performed within biological groups to avoid borrowing information across conditions and thereby attenuating true abundance differences. The benefits of this strategy were most evident in sparse quantitative regimes, such as low-input and single-cell DIA, where JUMPlion improved representation of low-abundance precursors and increased detection of differential protein abundance while maintaining low empirical differential-abundance FDRs.

Our results further indicate that complementary MS1- and MS2-derived evidence can improve DIA quantification, but that their relative value is platform dependent. Current DIA workflows are typically dominated by MS2 production measurements, whereas MS1 information is often secondary or not systematically incorporated into final quantitative estimates^17, 18^. Here, MS1-derived evidence provided clear benefits in timsTOF Pro 2 data, particularly in low-input settings, but yielded less consistent gains in the Exploris 480 and Orbitrap Astral datasets. These findings suggest that MS1 and MS2 should not be viewed as universally interchangeable or as competing alternatives. Rather, they represent complementary sources of quantitative information whose utility depends on acquisition-specific sampling density, signal quality, and interference structure. More broadly, these results support a more acquisition-aware view of DIA quantification, in which the balance between MS1- and MS2-derived evidence may need to be tuned to instrument platform and experimental context.

Several limitations should be considered. First, the datasets analyzed here consist of relatively controlled benchmark and application datasets of modest size. Although they are well suited for assessing quantitative completeness, fold-change accuracy, and differential abundance performance, they do not fully capture the heterogeneity, batch complexity, and broader dynamic range encountered in large cohort-scale biological or clinical studies. Second, although JUMPlion consistently improved completeness and differential abundance sensitivity, the current method depends on FDR-controlled precursor identifications as the starting point for signal extraction and does not replace the identification stage itself. Finally, as with other DIA quantification approaches, severe interference and highly unbalanced study designs may define operating regimes in which ion-level recovery is less effective or requires additional safeguards.

In summary, JUMPlion addresses a major source of DIA sparsity by recovering precursor-associated ion evidence before higher-level summarization and by integrating complementary MS1- and MS2-derived measurements in an acquisition-aware manner. Across benchmark, low-input, and single-cell datasets, this strategy improves quantitative completeness and enhances downstream differential abundance analysis, with the largest gains observed in settings where sparse ion observability most strongly constrains measurement. More broadly, these results suggest that incomplete DIA quantification should be viewed not only as a problem of missing entries in a protein matrix, but also as a problem of incomplete ion-level signal observability that can, at least in part, be addressed directly in raw data.

## Methods

### JUMPlion algorithm

An overview of the JUMPlion workflow is presented in **Fig. 1**. Detailed methods for each step are described below.

### Prediction of missing retention time and ion mobility values

Missing retention time (RT) and ion mobility (IM) values were predicted using run-specific calibration models constructed from high-confident precursors. By default, precursors detected in at least three runs were considered for model construction, and those showing high RT or IM variability across runs were excluded. The remaining precursors were ordered according to their library RT or IM values and used to build the calibration model. For each run, a piecewise spline regression model was fitted to map library RT (or IM) values to observed run-specific values. The resulting calibration function was then used to predict missing RT or IM values for precursors lacking observations in that run.

### Estimation of observable isotopic peaks

The number of observable isotopic peaks for each precursor was estimated using a Poisson approximation based on the averagine model. Specifically, the expected number of heavy-isotope substitutions arising from naturally occurring isotopes was assumed to scale linearly with peptide mass and was approximated as *λ* = 0.00052 × *m*, where *m* is the monoisotopic peptide mass in Daltons. The isotopic envelope was modeled using a Poisson distribution, and the most probable isotope peak (mode) was used as the reference for maximum theoretical intensity. Isotopic peaks were then evaluated sequentially beginning with the monoisotopic peak (M+0), and peaks with relative intensity greater than or equal to 10% of the maximum were considered observable. The total number of observable isotopic peaks was defined as the number of peaks from M+0 through the highest isotope peak satisfying this threshold.

### Determination of the representative extracted ion chromatogram

For each precursor, a representative extracted ion chromatogram (XIC) was selected from associated MS1- and/or MS2-level XICs to define the precursor elution profile. Candidate XICs were ranked using a composite score combining three metrics: (i) a correlation score reflecting similarity to other XICs assigned to the same precursor, (ii) a peak-shape score reflecting conformity to a Gaussian-like chromatographic profile with balanced signal around the apex, and (iii) an intensity score based on integrated signals. Individual metrics were normalized and combined using a weighted sum. For each highly ranked candidate XIC, the chromatographic profile was smoothed and the apex was refined by locating the local maximum intensity. Peak width was then estimated as the full width at half maximum (FWHM). Candidate XICs were required to satisfy predefined quality criteria, including a minimum peak-shape score and sufficient apex intensity. The highest-ranked XIC meeting these criteria was selected as the representative XIC for the precursor elution profile.

### Quantification of extracted ion chromatograms

Ion-level quantification was performed by integrating signal intensity within a chromatographic window centered on the apex of the representative precursor elution profile. The quantification window was defined using the defined apex and the estimated peak width, spanning the interval from (apex – FWHM) to (apex + FWHM). For each XIC, similarity to the representative elution profile was quantified using a correlation-based XIC-quality score, and the integrated signal within the quantification window was used as the raw quantity. Each XIC was then compared with the representative profile in the vicinity of the apex to estimate its expected signal intensity relative to the representative XIC. Observed intensities exceeding the profile-scaled expectation were truncated using separate scaling factors for MS1- and MS2-level XICs, after which XIC quantities were recalculated from the filtered XICs.

### Local inference of ion-level missingness

For each precursor, MS1- and MS2-level XIC quantities were processed separately across runs using a quality-aware inference procedure termed Local Inference of ION-level missingness using Neighborhood estimation (LION). XICs with low XIC quality or zero intensity were treated as missing, and XICs exhibiting excessive missingness across runs were excluded from downstream summarization. Remaining missing values were inferred separately for MS1- and MS2-level quantities by replacing missing entries with scaled estimates derived from neighborhood-specific minimum values computed among samples sharing the same experimental condition, as defined by sample metadata.

### Precursor-level summarization

For each precursor, MS1- and MS2-derived quantities were summarized independently from corresponding ion-level XIC measurements using a median-based summarization framework. Ion-level quantities were first log_2_-transformed, and the mean log_2_ intensity of each ion across runs was used to center that ion. Sample-specific medians of the centered values were then calculated to obtain precursor-level log_2_ intensity ratios. A representative abundance estimate was derived from the three most abundant ions and used to rescale these ratios. Final precursor-level quantities were obtained by back-transformation to linear intensity space.

### Protein-level summarization

Protein abundances were calculated from precursor-level quantities. Because each precursor could contribute both MS1- and MS2-derived measurements, JUMPlion supports three protein-level summarization modes: MS1-only, MS2-only, and integrated MS1/MS2 quantification. For protein groups supported by at least three qualified precursors, protein-level quantities were summarized using a median-based framework analogous to precursor-level summarization. For proteins supported by fewer than three qualified precursors, the precursor with the highest mean quality score was used as the representative estimate of protein abundance.

### Software implementation

JUMPlion was implemented in Python as a cross-platform software package supporting both graphical user interface (GUI) and command-line interface (CLI) execution. In CLI mode, user-defined parameters can be specified in a configuration file, enabling automated execution of the full quantification workflow.

### DIA-MS datasets

The HYE (human, yeast, and *E. coli*) peptide-mixture DIA benchmark datasets generated on timsTOF Pro 2 (loading amounts: 200 ng and 400 ng) and Exploris 480 (loading amount: 120 ng) instruments were downloaded from the MassIVE Consortium (MSV000092489)^28^. In the original study, samples were prepared using both a standard manual workflow and an automated OT-2 liquid-handling workflow. Because comparable numbers of proteins were quantified between the two preparation methods^28^, samples with the same loading amount were combined for downstream analysis. The Orbitrap Astral HYE DIA benchmark dataset (loading amount: 200 ng) was downloaded from the ProteomeXchange Consortium (PXD046444)^29^. The low-input HYE benchmark dataset generated on timsTOF Pro 2 (loading amount: 200 pg) and the single-cell DIA proteomics dataset generated on timsTOF Pro were downloaded from the ProteomeXchange Consortium via the iProX partner repository (PXD056832; IPX0009767000)^31^. For the low-input benchmark and single-cell datasets, DIA-NN (version 1.9.2) and Spectronaut (version 19.5.241126.62635) report files provided by the original study were also downloaded and used in downstream analyses.

### Protein sequence databases

Human (UP000005640; 20,647 entries), yeast (UP000002311; 6,065 entries), and *E. coli* (UP000000625; 4,402 entries) protein sequences were downloaded from UniProt release 2025_06. These FASTA files were combined for spectral-library generation and protein annotation.

### DIA-NN analysis

An in silico spectral library was generated in DIA-NN^17, 38^ (version 2.2.0 Academia) using the combined human, yeast, and *E. coli* FASTA database. Library-generation parameters were as follows: protease, Trypsin/P; maximum missed cleavages, 2; peptide length, 7–30 amino acids; fixed modification, carbamidomethylation of cysteine; variable modification, oxidation of methionine; protein N-terminal methionine excision enabled; precursor charge states, 2–4; precursor *m/z* range, 300–1400; and fragment *m/z* range, 100–1700. The resulting in silico library was used to analyze the timsTOF Pro 2, Exploris 480, and Orbitrap Astral HYE benchmark datasets^28, 29^. DIA-NN search and quantification settings were as follows: precursor FDR, 1%; mass accuracy for both MS1 and MS2 set to 15 ppm for timsTOF data and to automatic inference (0) for Exploris 480 and Orbitrap Astral data; scan window, automatic inference (0); match-between-runs enabled; unrelated runs disabled; protein inference enabled with proteotypicity set to “Genes”; scoring strategy set to “Peptidoforms”; machine-learning mode set to “NNs (cross-validated)”; quantification strategy set to QuantUMS (high precision); and cross-run normalization set to RT-dependent. The --export-quant option was used to export raw production quantities. The main report and empirical spectral library generated by DIA-NN were used as inputs for JUMPlion analysis. The ‘pr_matrix’ and ‘pg_matrix’ outputs were used as precursor- and protein-level quantification results, respectively, for downstream differential abundance analysis.

### Spectronaut analysis

DirectDIA analysis was performed in Spectronaut^2^ (version 20.3.251119.92449) using the combined human, yeast, and *E. coli* FASTA database. Default settings were used throughout, and the same analysis configuration was applied to benchmark datasets acquired on timsTOF Pro 2, Exploris 480, and Orbitrap Astral instruments. Both precursor-level and protein-level report files were exported for downstream comparison. For precursor-level quantification, the columns “EG.PrecursorId” and “EG.TotalQuantity (Settings)” were used. For protein-level quantification, the columns “PG.ProteinAccessions” and “PG.Quantity” were used.

### Definition of quantifiable proteins

Quantifiable proteins were defined using dataset-specific completeness criteria. For the timsTOF Pro 2 200 ng and 400 ng datasets and the Exploris 480 dataset^28^, proteins with valid intensity values in at least 5 of 8 replicates per sample group were considered quantifiable. For the Orbitrap Astral dataset^29^, proteins were required to be detected in all three replicates. For the low-input timsTOF Pro 2 HYE benchmark dataset^31^, proteins detected in at least 3 of 6 replicates were retained. For the single-cell proteomics dataset^31^, proteins with valid intensity values in at least 5 of 15 cells per treatment group were considered quantifiable. In all datasets, only proteins supported by at least two precursors were retained for downstream analysis.

### Differential abundance analysis and performance evaluation

The protein-level quantification matrices outputted by JUMPlion, DIA-NN, and Spectronaut were log_2_-transformed. Differential abundance analysis between groups was then performed using limma^33^ (version 3.62.2) through a customized implementation of the LFQ_bout R script^30^ (https://github.com/t-jumel/LFQb) in R (version 4.4.3). For all benchmark datasets, the smallest expected absolute log_2_ fold change (log_2_FC) between conditions was 1. Thus, proteins with |log_2_FC| > 0.5 and Benjamini–Hochberg adjusted *P* < 0.05 were classified as differentially abundant proteins (DAPs). For benchmark-based performance evaluation, ground truth was defined according to the known species mixing ratios in the HYE samples. Human proteins were expected to remain unchanged, yeast proteins were expected to increase in abundance, and *E. coli* proteins were expected to decrease in abundance. Proteins meeting the differential abundance criteria in the expected direction were classified as true positives (TPs). Proteins meeting the criteria in the opposite direction or among expected unchanged proteins were classified as false positives (FPs). Proteins from expected changing groups that failed to meet the criteria were classified as false negatives (FNs), and unchanged human proteins that did not meet the criteria were classified as true negatives (TNs). Receiver operating characteristic (ROC) curves were generated using scikit-learn (version 1.6.1) by plotting true positive rate (TPR) against false positive rate (FPR) on varying adjusted *P*-value thresholds. Partial area under the ROC curve (pAUC) was calculated within the range FPR ≤ 0.1.

### Single-cell data analysis

Single-cell protein quantification matrices were log_2_-transformed and normalized using SCnorm^32^, followed by batch correction with limma^33^ (removeBatchEffect). Treatment group was included during batch correction to preserve condition-associated abundance differences. Gene set enrichment analysis (GSEA)^34^ was performed on protein-level log_2_FCs using the R package genekitr^39^ (version 1.2.8) in R (version 4.4.3), with Gene Ontology Biological Process^35^ and Reactome^36^ gene sets used as reference collections. Gene sets with adjusted *P* < 0.05 were considered significantly enriched.

## Code availability

JUMPlion source code and documentation are available at https://github.com/yingxue-fu/jumplion under the Apache 2.0 license.

## Acknowledgements

This work was supported by NIH grants AG092468, AG064909 and AG069701, and the American Lebanese Syrian Associated Charities (ALSAC) foundation. We gratefully thank Jay M. Yarbro for testing JUMPlion.

## Author contributions

Conceptualization: Y.Fu, Z.Yuan, X.Wang, J.Peng, A.High. Data curation: Y.Fu, Z. Yuan, S.Byrum, L.Wu. Formal analysis: Y.Fu, Z. Yuan, S.Byrum, L.Wu, X.Wang. Funding Acquisition: J.Peng, A.High. Investigation: Y.Fu, Z. Yuan, S.Byrum, L.Wu, X.Wang. Methodology: Y.Fu, Z.Yuan, X.Wang, J.Peng. Software: Y.Fu. Resources: A.High. Supervision: Z.Yuan, X.Wang, A.High. Visualization: Y.Fu. Writing-original draft: Y.Fu. Writing-review and editing: All authors.

## Competing interests

The authors declare no competing interests.

## Supplementary Figures

**Supplementary Fig. 1.**
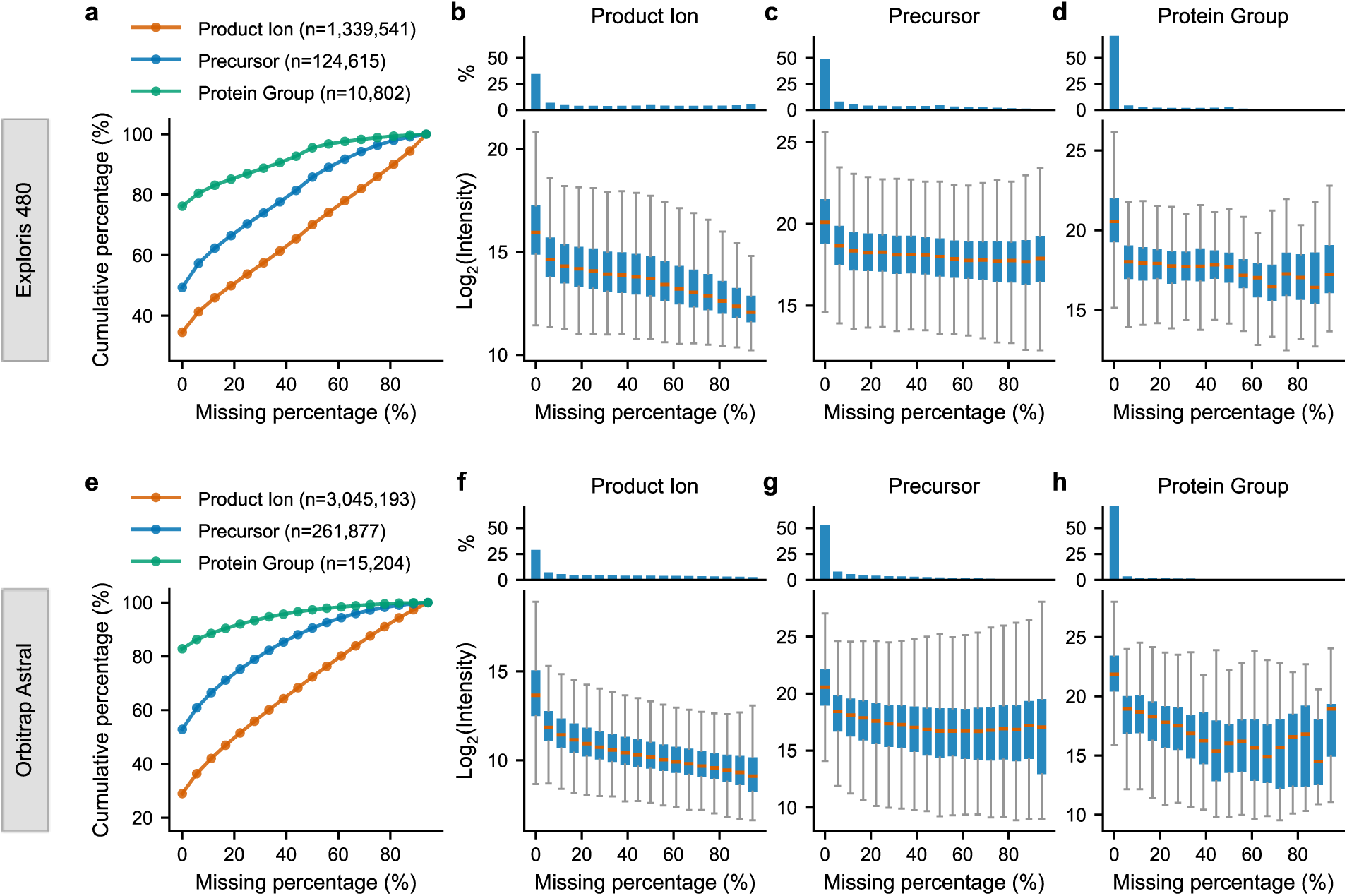
The extent of missingness at different quantification levels in Exploris 480 and Orbitrap Astral benchmark datasets. **a–d**, Exploris 480 dataset. **a**, Cumulative percentages of product ions, precursors, and protein groups on different levels of missingness. **b–d**, Relationship between missing-value percentage and log_2_ intensity at the production (**b**), precursor (**c**), and protein-group (**d**) levels. **e–h**, Same as **a–d** for the Orbitrap Astral dataset.

**Supplementary Fig. 2.**
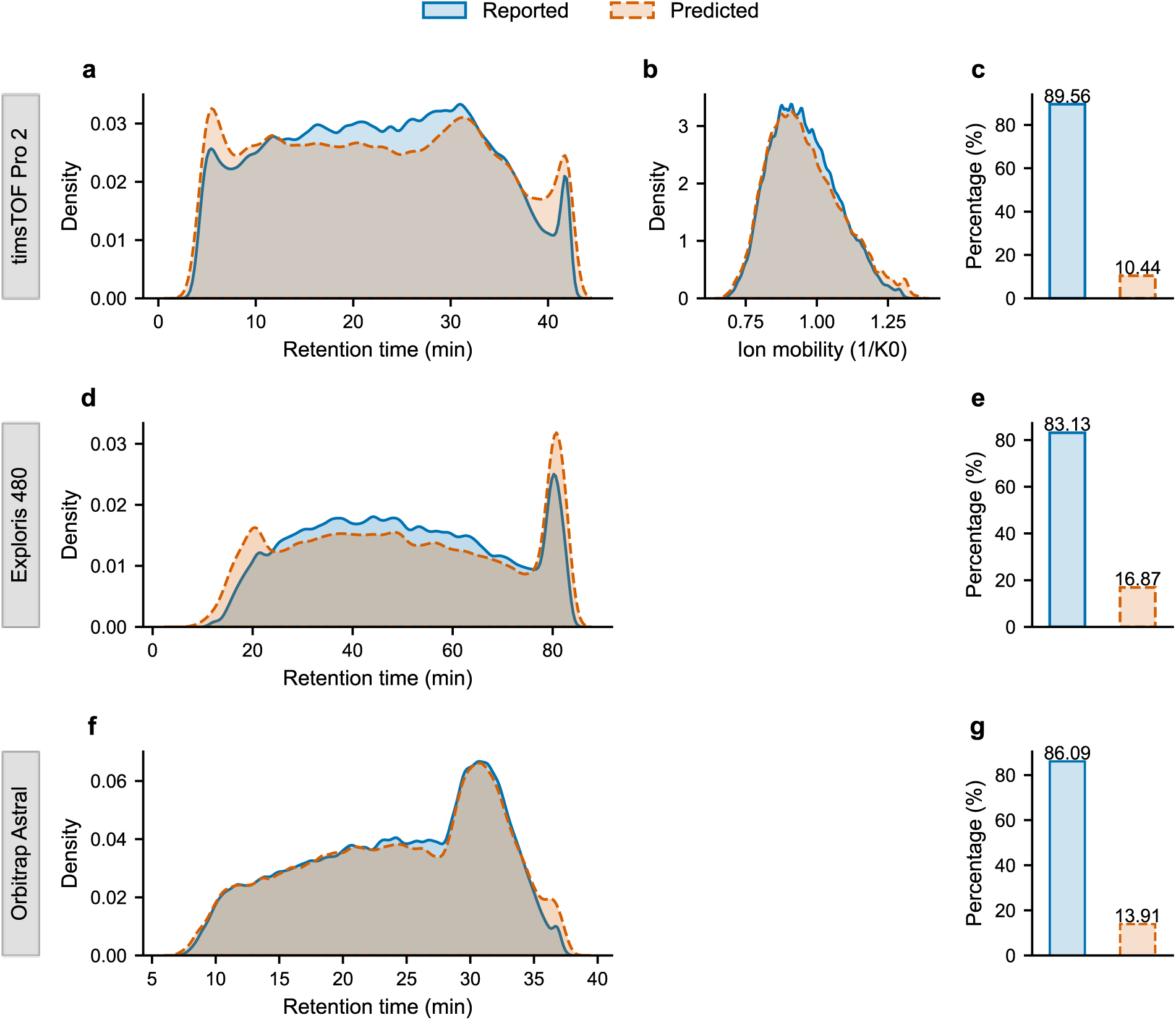
Reported and inferred chromatographic coordinates used for signal extraction. **a–c**, timsTOF Pro 2 dataset. **a**, Distributions of reported and JUMPlion-predicted retention times (RTs). **b**, Distributions of reported and JUMPlion-predicted ion mobility (IM) values. **c**, Proportions of reported and JUMPlion-predicted RT and IM values. **d,e**, Same as **a** and **c**, respectively, for the Exploris 480 dataset. **f,g**, Same as **a** and **c**, respectively, for the Orbitrap Astral dataset.

**Supplementary Fig. 3.**
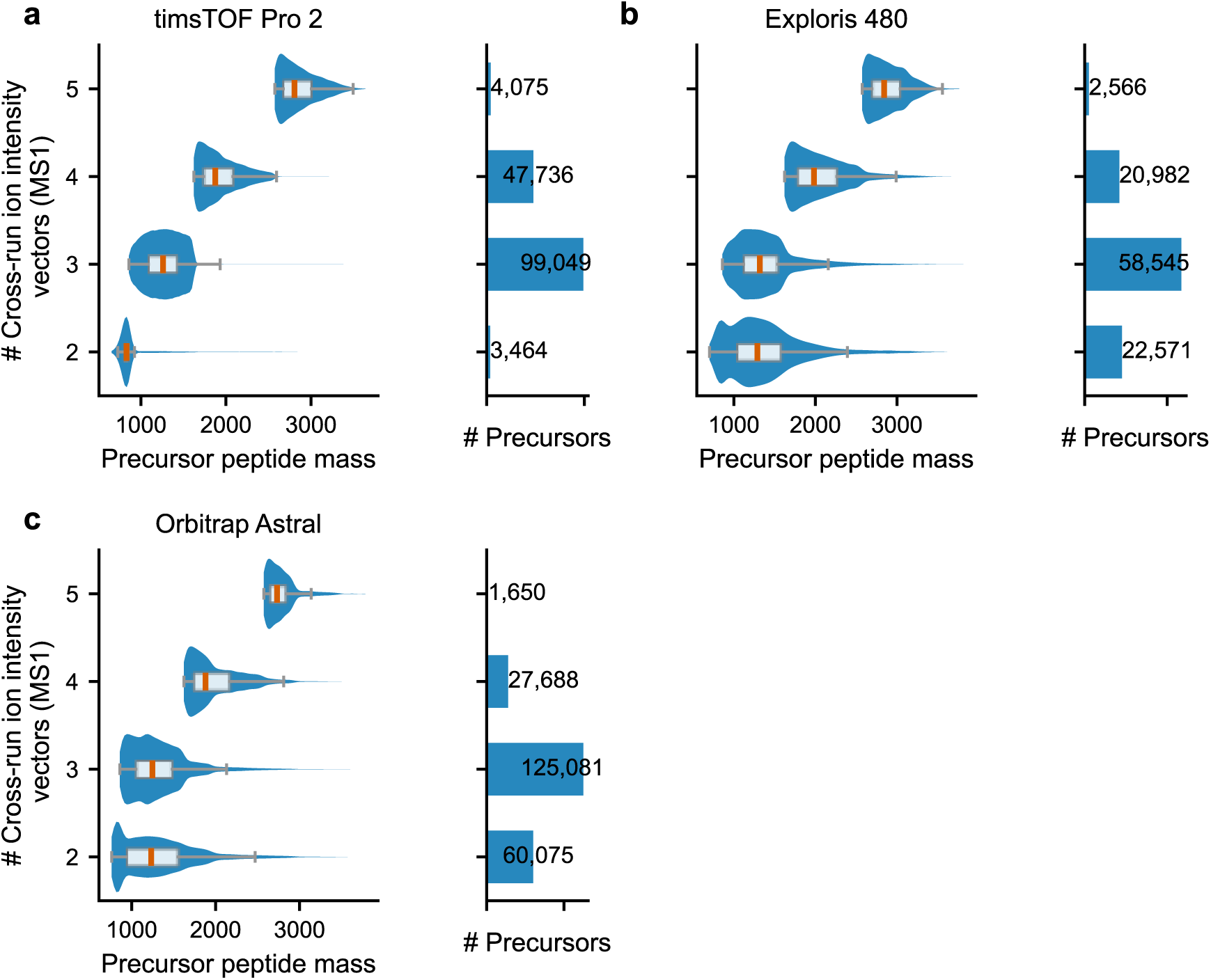
Relationship between precursor peptide mass and the number of MS1-level cross-run ion intensity vectors. **a**, timsTOF Pro 2 dataset. **b**, Exploris 480 dataset. **c**, Orbitrap Astral dataset.

**Supplementary Fig. 4.**
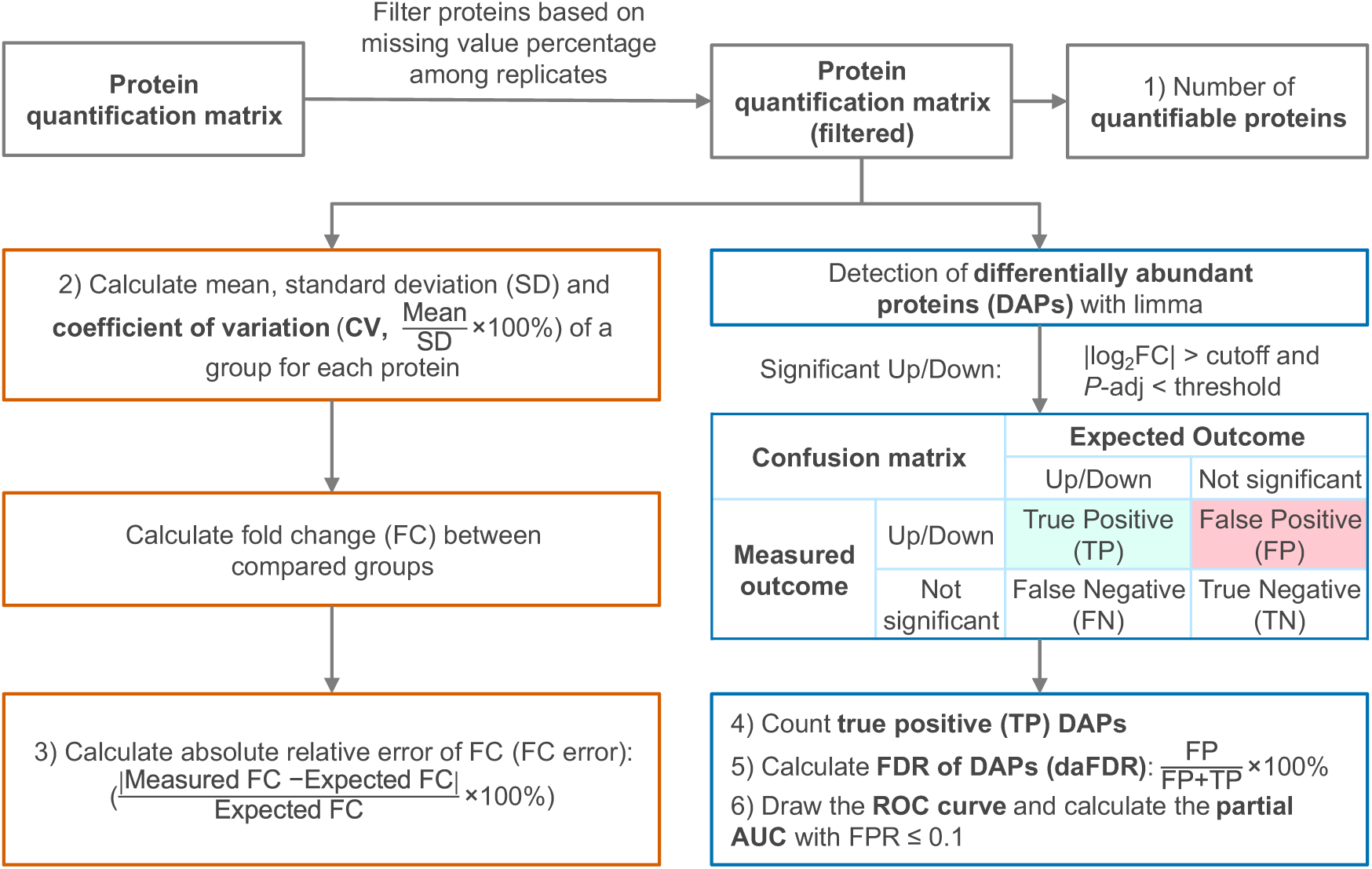
The metrics used for evaluating the quantification performance.

**Supplementary Fig. 5.**
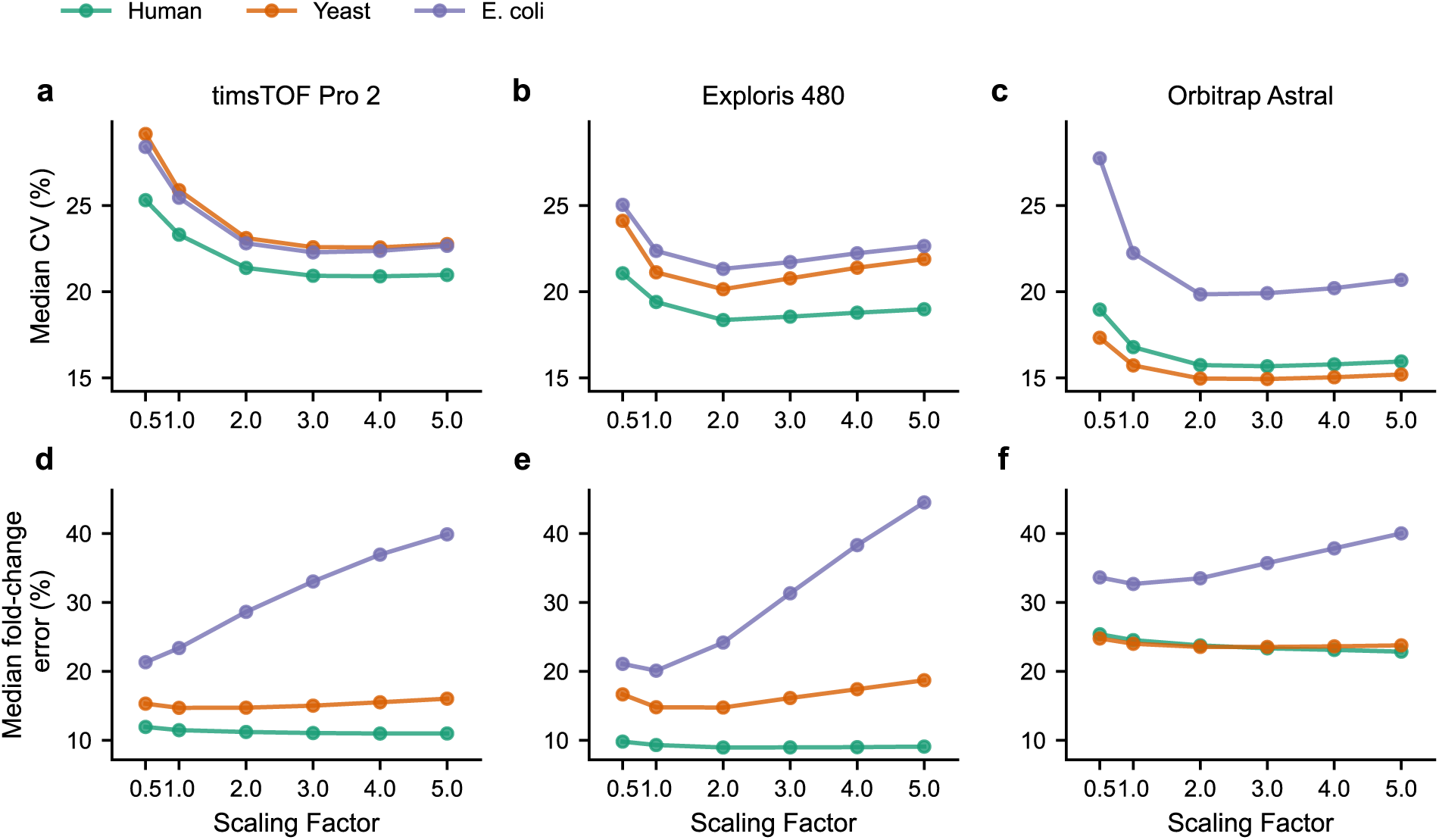
Effect of the local-minimum scaling factor on precursor-level quantification. **a–c**, Median coefficient of variation (CV) as a function of the scaling factor applied to the local minimum intensity for the timsTOF Pro 2 (**a**), Exploris 480 (**b**), and Orbitrap Astral (**c**) datasets. **d–f**, Median fold-change error for the corresponding datasets.

**Supplementary Fig. 6.**
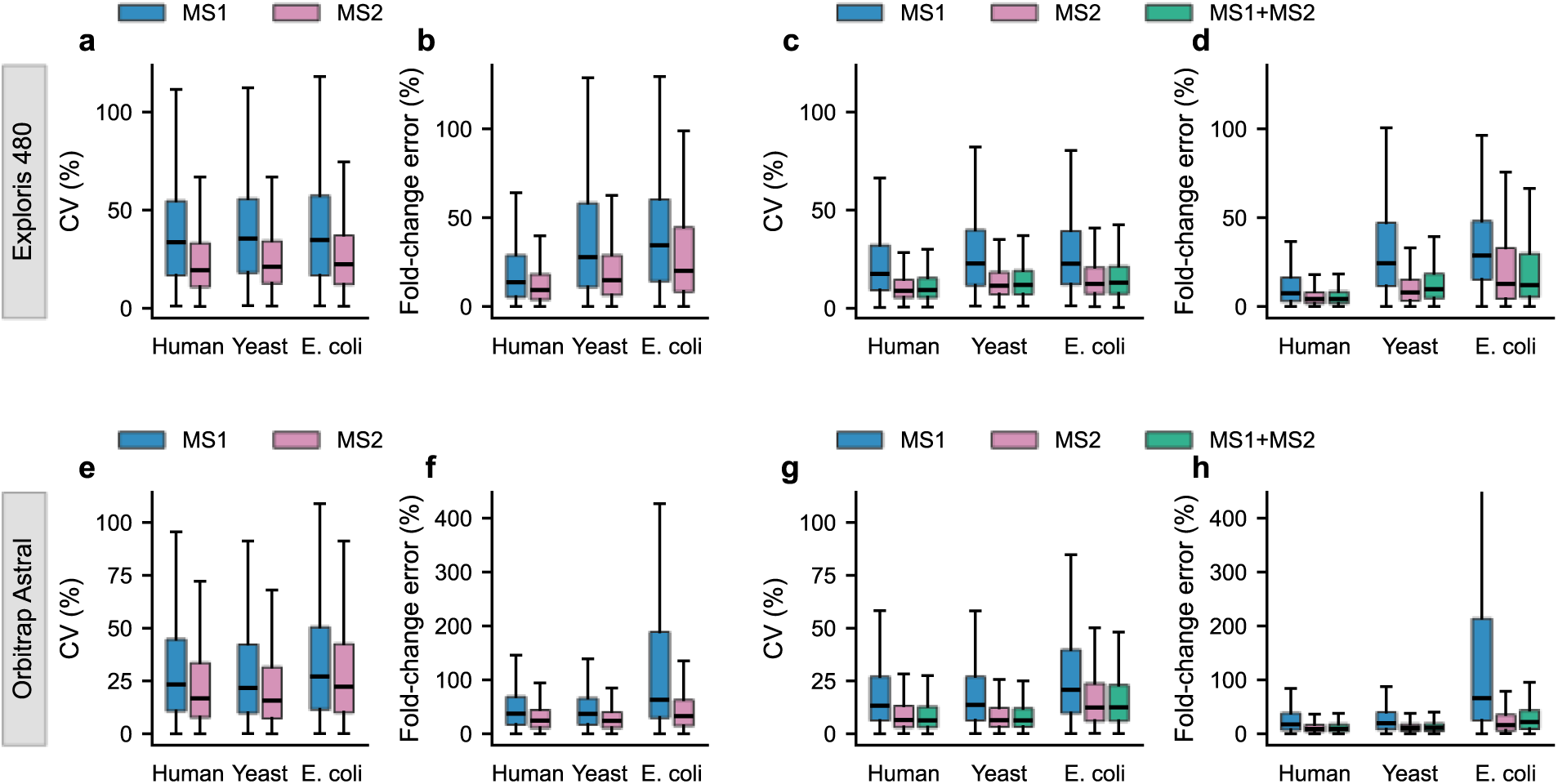
Precursor- and protein-level quantification performance in additional DIA platforms. **a–d**, Exploris 480 dataset. **a,b**, CV (**a**) and fold-change error (**b**) of precursor quantities summarized using LION-inferred MS1- and MS2-level ion intensities. **c,d**, CV (**c**) and fold-change error (**d**) of protein quantities summarized using MS1-only, MS2-only, and integrated MS1/MS2 strategies. **e–h**, Same as **a–d** for the Orbitrap Astral dataset.

**Supplementary Fig. 7.**
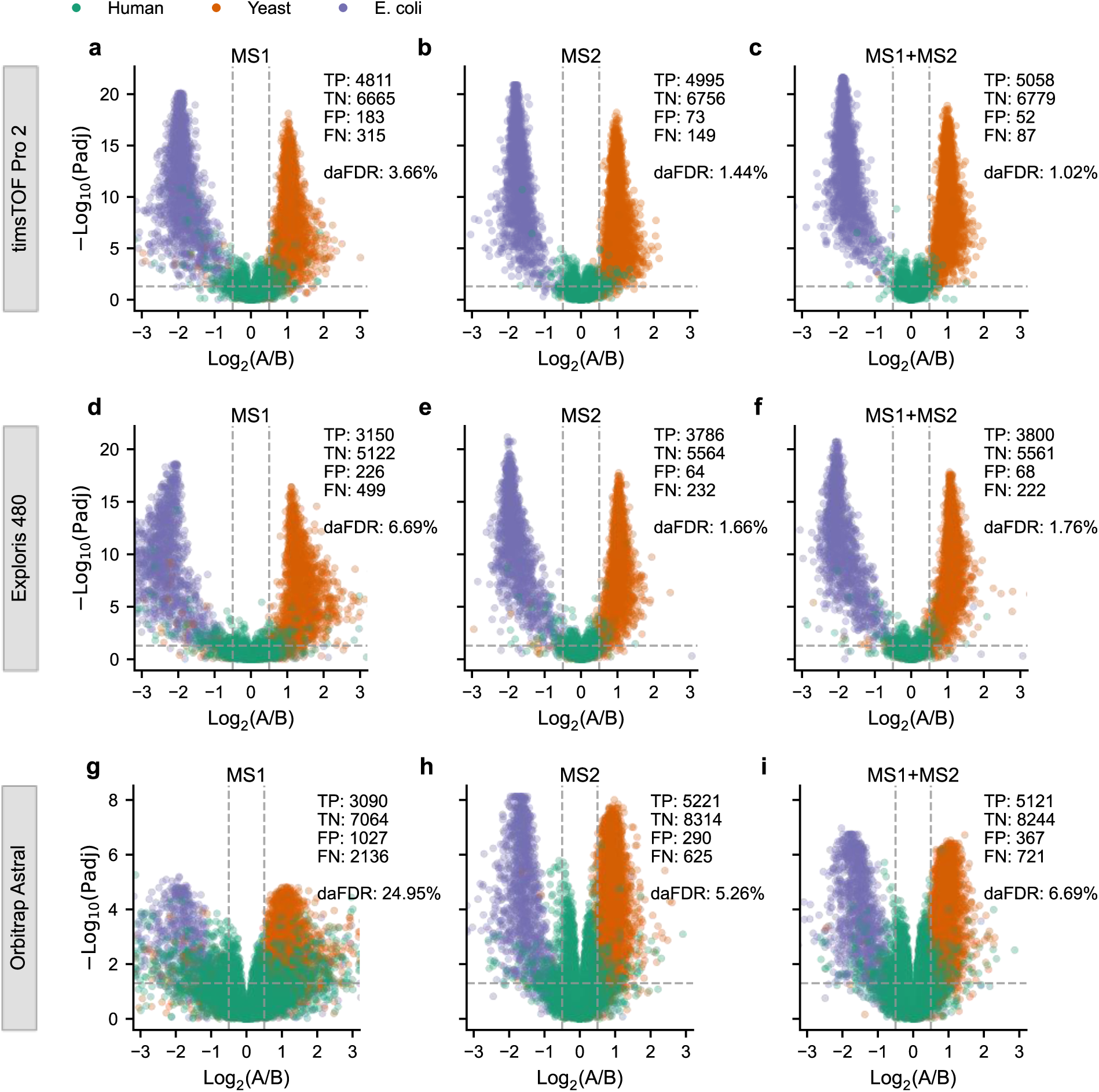
Differential abundance analyses for the protein quantities summarized using MS1-only, MS2-only and integrated MS1/MS2 approaches. **a–c**, timsTOF Pro 2 dataset. Volcano plots of differential protein abundance analysis using protein quantities summarized using MS1-only (**a**), MS2-only (**b**) and integrated MS1/MS2 (**c**) approaches. Points are colored by protein species of origin. The numbers of true positives (TP), true negatives (TN), false positives (FP) and false negatives (FN), along with the differential abundance false discovery rate (daFDR; FP/(TP+FP)), are indicated. **d–f**, same as **a–c** but for the Exploris 480 dataset. **g–i**, same as **a–c** but for the Orbitrap Astral dataset.

**Supplementary Fig. 8.**
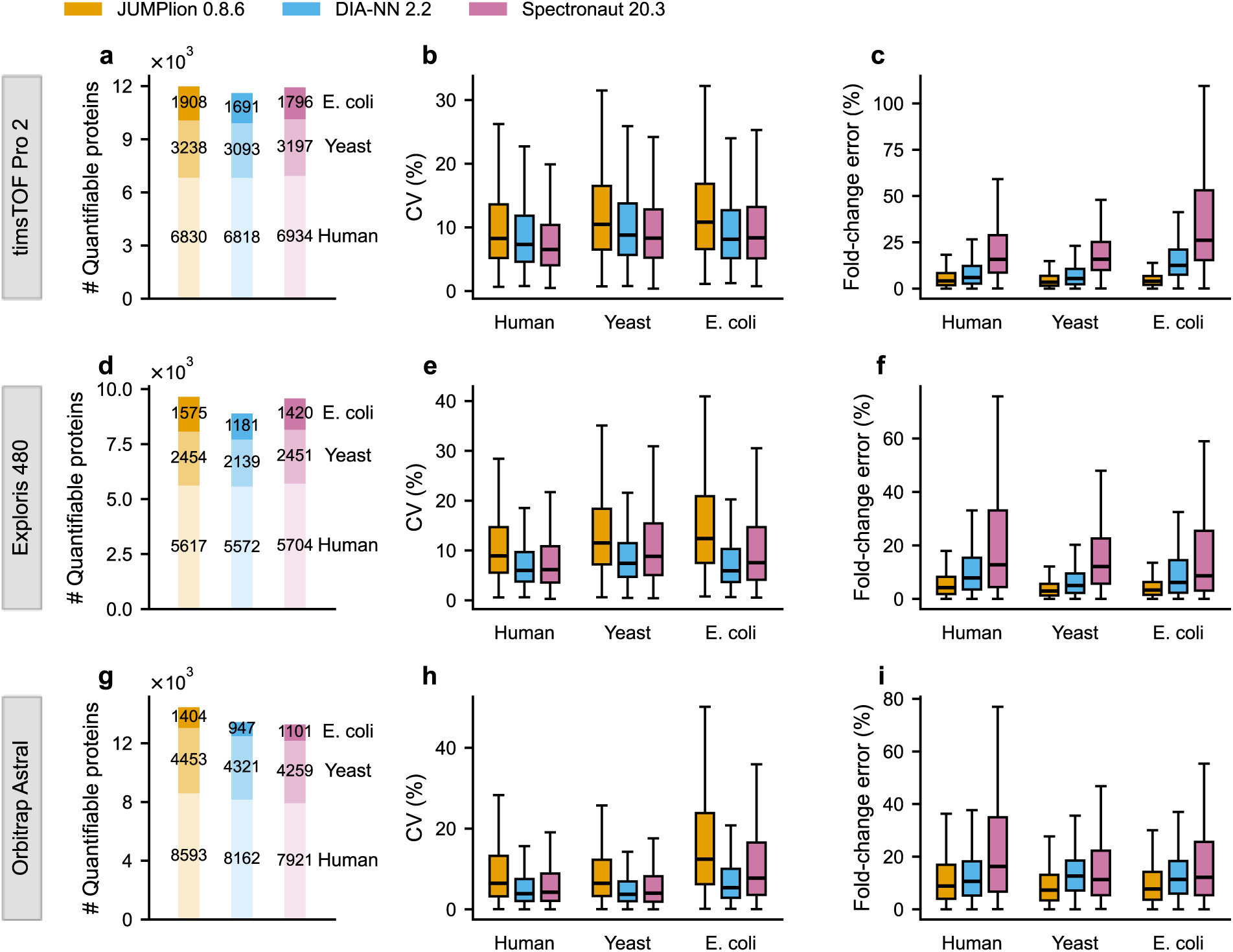
Protein-level quantification performance in JUMPlion v0.8.6, DIA-NN v2.2 and Spectronaut v20.3. **a–c**, timsTOF Pro 2 dataset. Number of quantifiable proteins (**a**), CV (**b**) and fold-change error (**c**) based on protein quantities derived from JUMPlion v0.8.6, DIA-NN v2.2 and Spectronaut v20.3. **d–f**, same as **a–c** but for the Exploris 480 dataset. **g–i**, same as **a–c** but for the Orbitrap Astral dataset.

**Supplementary Fig. 9.**
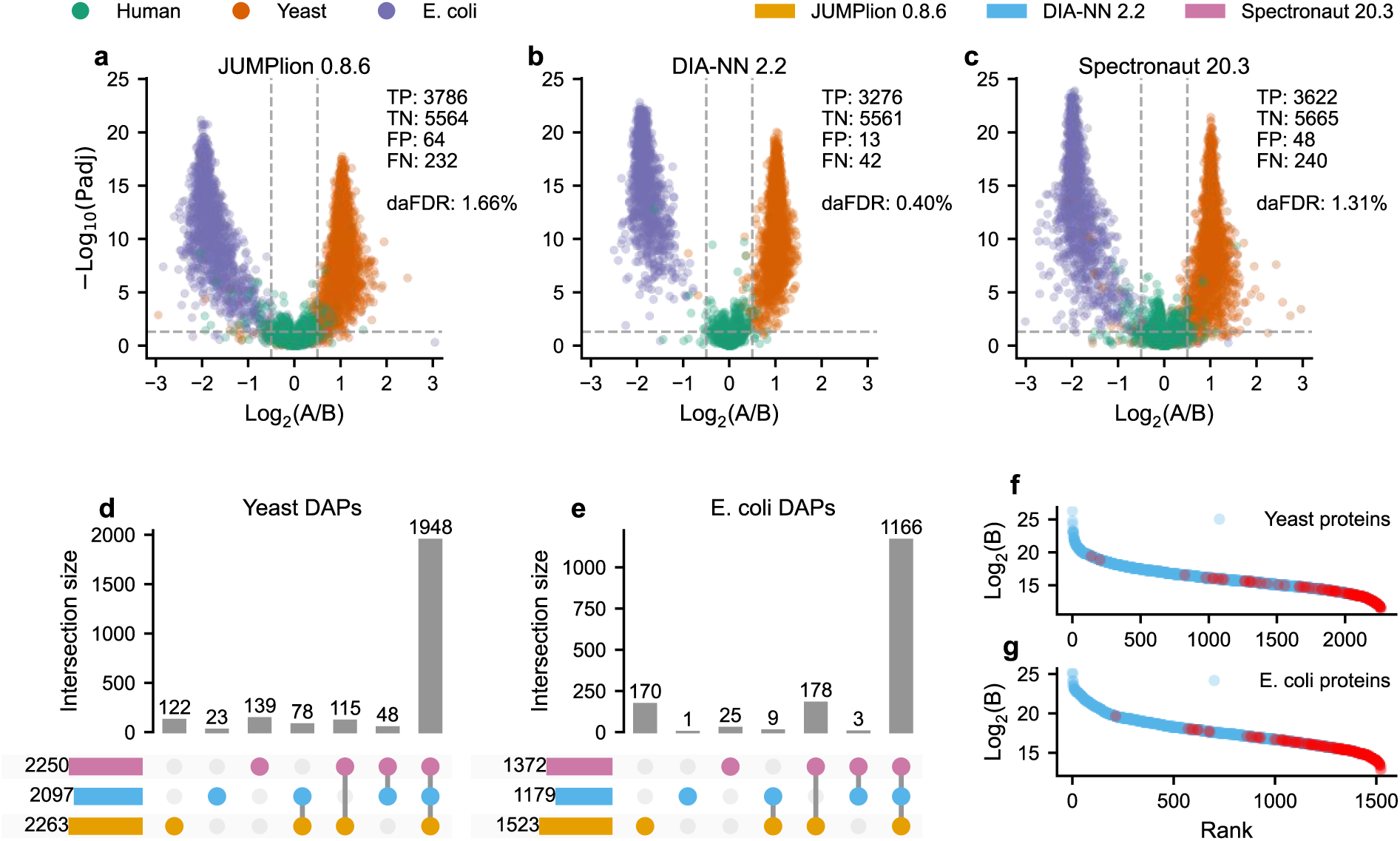
Differential abundance analyses for the Exploris 480 dataset. **a–c**, Volcano plots of differential protein abundance analysis using protein quantities derived from JUMPlion v0.8.6 (**a**), DIA-NN v2.2 (**b**) and Spectronaut v20.3 (**c**). Points are colored by protein species of origin. The numbers of TP, TN, FP and FN, along with the daFDR (FP/(TP+FP)), are indicated. **d,e**, UpSet plots showing unique and shared differentially abundant proteins (DAPs) from yeast (d) and *E. coli* (e) identified by JUMPlion, DIA-NN and Spectronaut. **f,g**, Ranked log₂ intensities of all yeast and *E. coli* proteins, with DAPs uniquely identified using JUMPlion-derived quantities highlighted in red.

**Supplementary Fig. 10.**
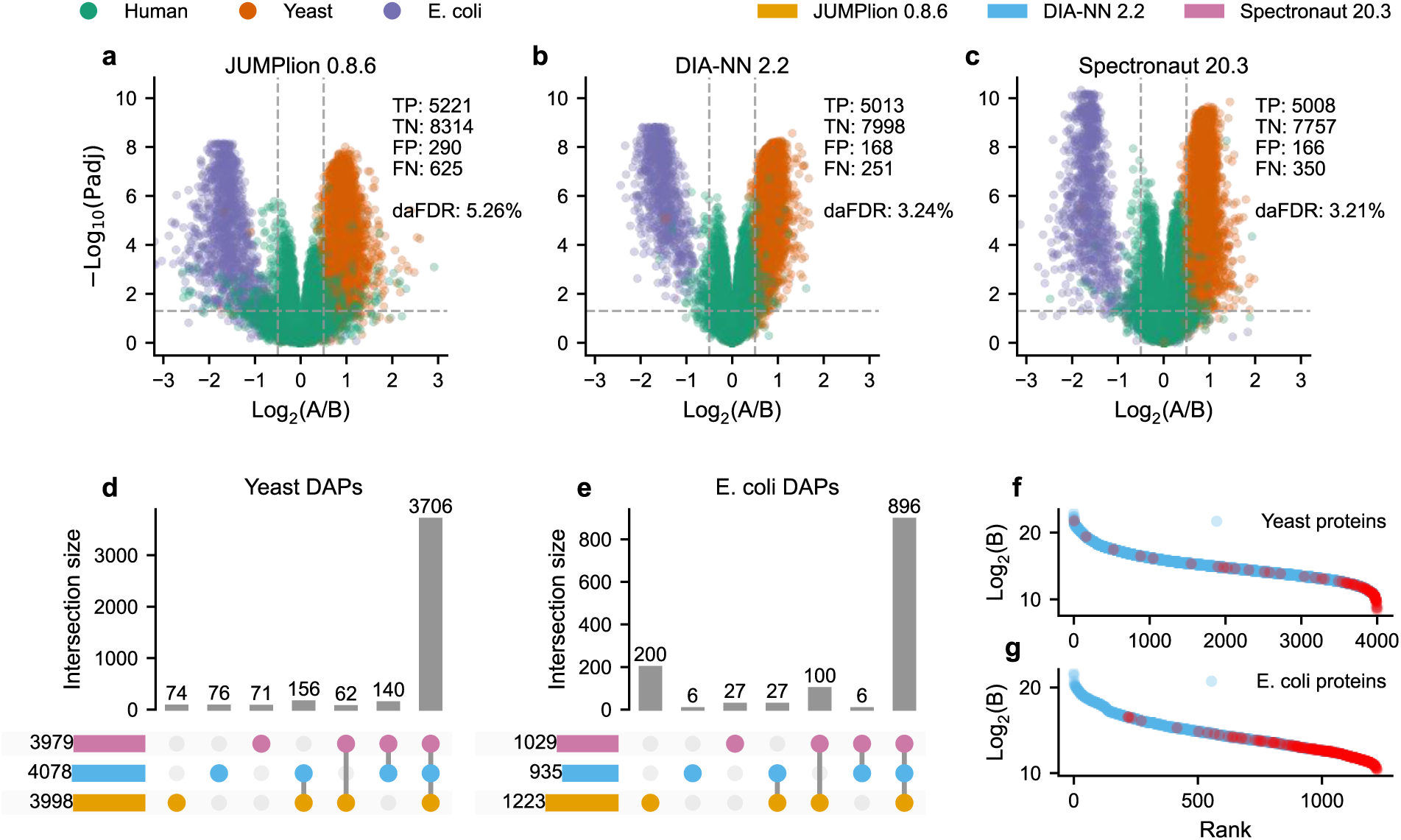
Differential abundance analyses for the Orbitrap Astral dataset. **a–c**, Volcano plots of differential protein abundance analysis using protein quantities derived from JUMPlion v0.8.6 (a), DIA-NN v2.2 (b) and Spectronaut v20.3 (c). Points are colored by protein species of origin. The numbers of TP, TN, FP and FN, along with the daFDR (FP/(TP+FP)), are indicated. **d,e**, UpSet plots showing unique and shared DAPs from yeast (d) and *E. coli* (e) identified by JUMPlion, DIA-NN and Spectronaut. **f,g**, Ranked log₂ intensities of all yeast and *E. coli* proteins, with DAPs uniquely identified using JUMPlion-derived quantities highlighted in red.

**Supplementary Fig. 11.**
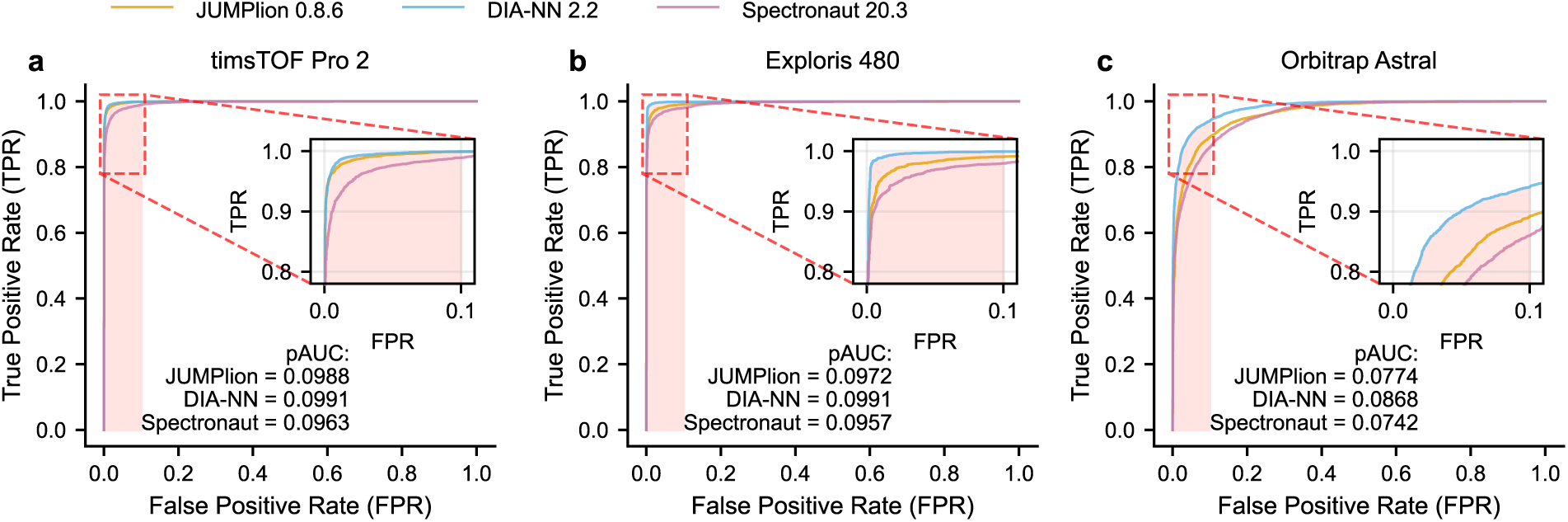
Receiver operating characteristic (ROC) curves for differential abundance benchmarking. ROC curves for timsTOF Pro 2 (**a**), Exploris 480 (**b**) and Orbitrap Astral (**c**) datasets ere generated using −log_10_(adjusted P) as the ranking score. The partial area under the curve (pAUC) within false positive rate (FPR) ≤ 0.1 is highlighted in light red.

**Supplementary Fig. 12.**
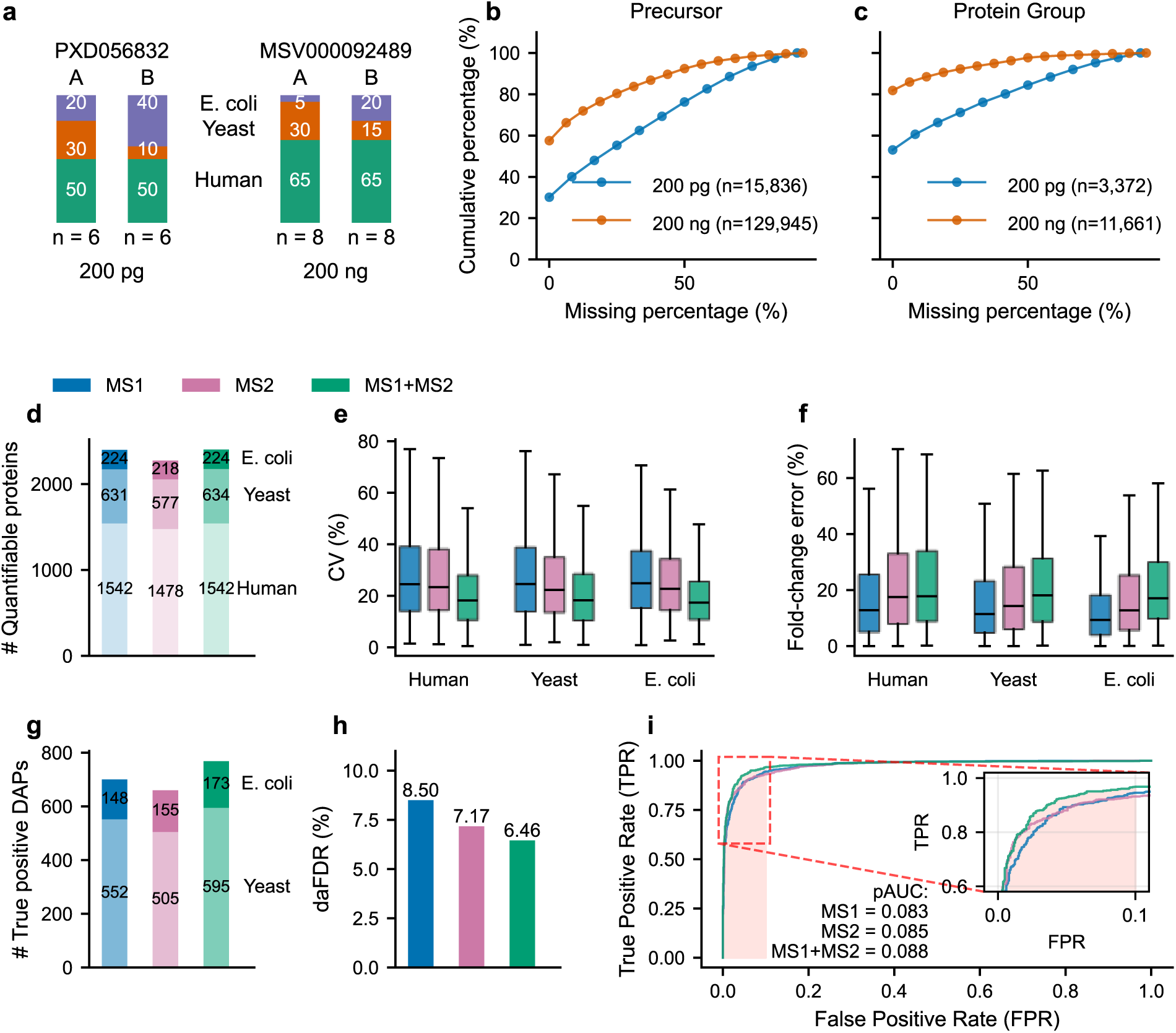
JUMPlion performance in low-input timsTOF Pro 2 DIA benchmarks. **a**, Overview of two timsTOF Pro 2 benchmark datasets differing by approximately three orders of magnitude in peptide loading amount (200 pg versus 200 ng). **b,c**, Cumulative percentages of precursors (**b**) and protein groups (**c**) on different levels of missingness in the 200 pg and 200 ng datasets. **d–i**, Quantification and differential abundance benchmarking for JUMPlion in the 200 pg dataset. **d**, Number of quantifiable proteins. **e**, CV. **f**, Fold-change error. **g**, Number of true positive DAPs. **h**, daFDR. **i**, ROC curves based on protein quantities summarized using MS1-only, MS2-only, and integrated MS1/MS2 strategies.

**Supplementary Fig. 13.**
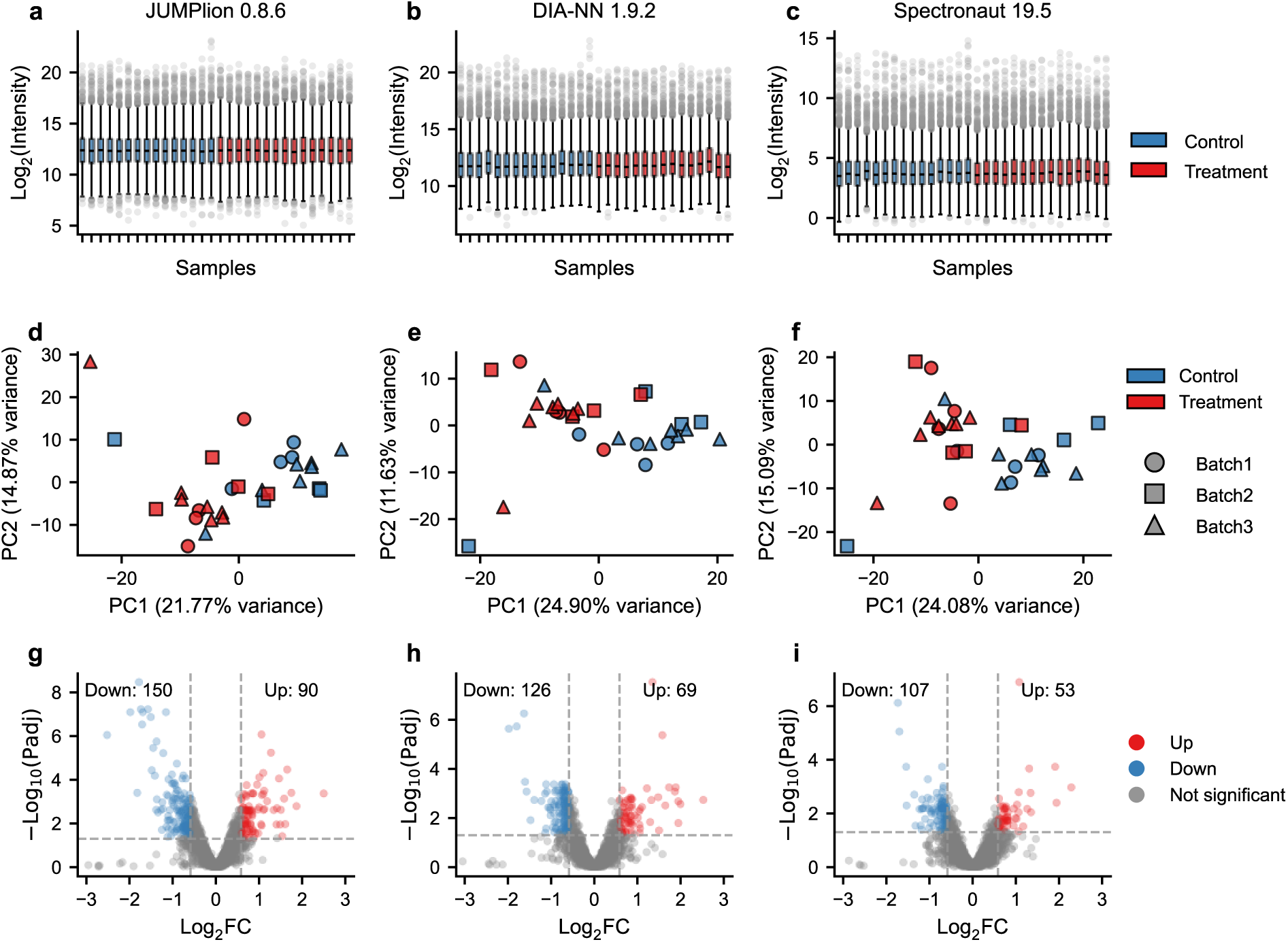
Normalization, batch correction, and differential abundance analysis of single-cell DIA data. **a–c**, Box plots of log_2_ protein intensities derived from JUMPlion (**a**), DIA-NN (**b**), and Spectronaut (**c**) after SCnorm normalization and limma-based batch correction. **d–f**, Principal component analysis plots generated using the top 500 most variable proteins in each quantification matrix. **g–i**, Volcano plots of differential protein abundance analysis using protein quantities derived from JUMPlion (**g**), DIA-NN (**h**), and Spectronaut (**i**). Points are colored as up-regulated, down-regulated, or not significant. The numbers of up- and down-regulated proteins are indicated.

**Supplementary Table 1.**
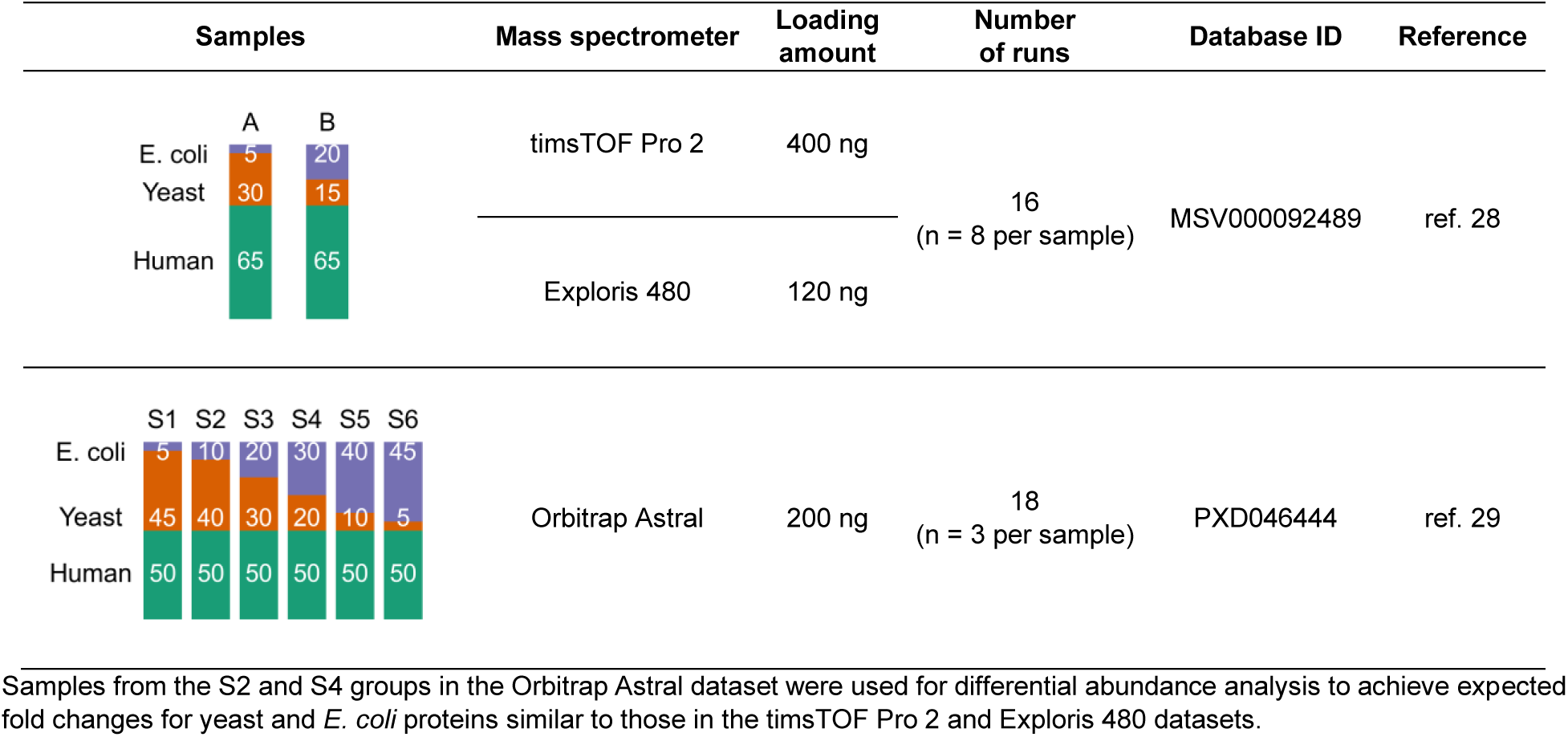
Benchmark DIA-MS datasets used in cross-platform analyses.

## Notes

### Competing Interest Statement

The authors have declared no competing interest.

